# Transethnic meta-analysis of genome-wide association studies identifies three new loci and characterizes population-specific differences for coronary artery disease

**DOI:** 10.1101/739979

**Authors:** Hiroshi Matsunaga, Kaoru Ito, Masato Akiyama, Atsushi Takahashi, Satoshi Koyama, Seitaro Nomura, Hirotaka Ieki, Kouichi Ozaki, Yoshihiro Onouchi, Shinichiro Suna, Soichi Ogishima, Masayuki Yamamoto, Atsushi Hozawa, Mamoru Satoh, Makoto Sasaki, Taiki Yamaji, Norie Sawada, Motoki Iwasaki, Shoichiro Tsugane, Keitaro Tanaka, Kokichi Arisawa, Hiroaki Ikezaki, Naoyuki Takashima, Mariko Naito, Kenji Wakai, Hideo Tanaka, Yasuhiko Sakata, Hiroyuki Morita, Yasushi Sakata, Koichi Matsuda, Yoshinori Murakami, Hiroshi Akazawa, Michiaki Kubo, Yoichiro Kamatani, Issei Komuro

**Affiliations:** From Laboratory for Cardiovascular Genomics and Informatics, RIKEN Center for Integrative Medical Sciences, Kanagawa, Japan; Department of Cardiovascular Medicine, Graduate School of Medicine, The University of Tokyo, Tokyo, Japan; Laboratory for Statistical Analysis, RIKEN Center for Integrative Medical Sciences, Kanagawa, Japan; Department of Genomic Medicine, Research Institute, National Cerebral and Cardiovascular Center, Osaka, Japan; Genome Science Division, Research Center for Advanced Science and Technologies, The University of Tokyo, Tokyo, Japan; Division for Genomic Medicine, Medical Genome Center, National Center for Geriatrics and Gerontology, Obu, Japan; Department of Public Health, Chiba University Graduate School of Medicine, Chiba, Japan; Department of Cardiovascular Medicine, Osaka University Graduate School of Medicine, Suita, Japan; Tohoku Medical Megabank Organization, Tohoku University, Sendai, Japan; Department of Preventive Medicine and Epidemiology, Tohoku Medical Megabank Organization, Tohoku University, Sendai, Japan; Iwate Tohoku Medical Megabank Organization, Iwate Medical University, Iwate, Japan; Division of Epidemiology, Center for Public Health Sciences, National Cancer Center, Tokyo, Japan; Center for Public Health Sciences, National Cancer Center, Tokyo, Japan; Department of Preventive Medicine, Faculty of Medicine, Saga, University, Saga, Japan; Department of Preventive Medicine, Institute of Biomedical Sciences, Tokushima University Graduate School, Tokushima, Japan; Department of Environmental Medicine and Infectious Diseases, Graduate School of Medical Sciences, Kyushu University, Fukuoka, Japan; Department of Public Health, Shiga University of Medical Science, Otsu, Japan; Department of Oral Epidemiology, Graduate School of Biomedical and Health Sciences, Hiroshima University; Department of Preventive Medicine, Nagoya University Graduate School of Medicine, Nagoya, Japan; Division of Epidemiology and Prevention, Aichi Cancer Center Research Institute, Nagoya, Japan; Department of Epidemiology, Nagoya University Graduate School of Medicine, Nagoya, Japan; Department of Cardiovascular Medicine, Tohoku University Graduate School of Medicine, Sendai, Japan; Department of Computational Biology and Medical Science, Graduate school of Frontier Sciences, The University of Tokyo, Tokyo, Japan; Division of Molecular Pathology, Institute of Medical Science, The University of Tokyo, Tokyo, Japan; RIKEN Center for Integrative Medical Sciences, Kanagawa, Japan; Kyoto-McGill International Collaborative School in Genomic Medicine, Kyoto University Graduate School of Medicine, Kyoto, Japan

**Keywords:** Coronary artery disease, Genome-wide association study, Japanese, Genetics, Single nucleotide polymorphism

## Abstract

**Background:** Genome-wide association studies (GWAS) provided many biological insights into coronary artery disease (CAD), but these studies were mainly performed in Europeans. GWAS in diverse populations have the potential to advance our understanding of CAD.

**Methods and Results:** We conducted two GWAS for CAD in the Japanese population, which included 12,494 cases and 28,879 controls, and 2,808 cases and 7,261 controls, respectively. Then, we performed transethnic meta-analysis using the results of the CARDIoGRAMplusC4D 1000 Genomes meta-analysis with UK Biobank. We identified 3 new loci on chromosome 1q21 (*CTSS*), 10q26 (*WDR11-FGFR2*), and 11q22 (*RDX*-*FDX1*). Quantitative trait locus analyses suggested the association of *CTSS* and *RDX*-*FDX1* with atherosclerotic immune cells. Tissue/cell type enrichment analysis showed the involvement of arteries, adrenal glands and fat tissues in the development of CAD. Finally, we performed tissue/cell type enrichment analysis using East Asian-frequent and European-frequent variants according to the risk allele frequencies, and identified significant enrichment of adrenal glands in the East Asian-frequent group while the enrichment of arteries and fat tissues was found in the European-frequent group. These findings indicate biological differences in CAD susceptibility between Japanese and Europeans.

**Conclusions:** We identified 3 new loci for CAD and highlighted the genetic differences between the Japanese and European populations. Moreover, our transethnic analyses showed both shared and unique genetic architectures between the Japanese and Europeans. While most of the underlying genetic bases for CAD are shared, further analyses in diverse populations will be needed to elucidate variations fully.

## Introduction

Coronary artery disease (CAD) remains a leading cause of death worldwide, despite recent improvements in its treatment and prevention. To date, genome-wide association studies (GWAS) have revealed more than 150 CAD loci and some of the CAD loci associated with conventional atherosclerotic risk factors.^1–6^ Especially, loci related to hypertension and dyslipidemia are major contributors to CAD. GWAS have also offered new insights into CAD by uncovering shared genetic mechanisms with other diseases, such as migraine headaches and autoimmune diseases.^7^ However, the currently known loci identified by GWAS explain only a small percentage (∼15%) of disease heritability,^6^ while a study of twins estimated a heritability of ∼50% for CAD.^8^ This strongly suggests that there still remain undetected loci associated with CAD.

CAD risk loci have been identified primarily in European populations. Because ethnicity and population variations can affect the genetic traits of patients, it is important to investigate genetic diversity in different populations to determine the genetic basis of a disease. Actually, the incidence, prevalence, and mortality of CAD vary across populations. The age-adjusted incidence rate (per 100,000 persons per year) for myocardial infarction (MI) in Japan increased from 7.4 in 1979 to 27 in 2008,^9^ while that in European countries ranged from 44 to 142, respectively.^10^ As a result of the westernization of Japanese food habits and the world’s most aged society, the number of Japanese patients diagnosed with CAD is rapidly increasing. Therefore, the importance of prediction, prevention and treatment for CAD is growing and studies for Japanese are needed to accumulate population-specific evidences to support it.

Herein, we present a meta-analysis of 2 Japanese GWAS for MIs, which includes 15,302 cases and 36,140 controls. We also present a trans-ethnic meta-analysis using the 2 Japanese GWAS datasets and the CARDIoGRAMplusC4D 1000 Genomes meta-analysis with UK Biobank data, which identified 3 novel loci. We then explore the pathophysiological significance of these novel loci and examine the differences in CAD-susceptibility loci between Japanese and Europeans.

## Methods

### Study design and samples in the Japanese GWAS

We performed two Japanese GWAS using independent case-control cohorts separately, since the two cohorts that encompassed cases for the first Japanese GWAS and the second Japanese GWAS were independent and unrelated to each other. We then conducted a meta-analysis of these GWAS (Figure 1). In the first Japanese GWAS, all cases of MI were selected from Biobank Japan (BBJ), which is a registry that collected DNA and serum samples at 12 cooperative medical institutions in Japan from June 2003 to March 2008. All control samples were obtained from Tohoku Medical Megabank Organization (ToMMo), Iwate Tohoku Medical Megabank Organization (IMM), the Japan Public Health Center-based Prospective (JPHC) Study,^11^ and the Japan Multi-Institutional Collaborative Cohort (J-MICC) Study. In the second Japanese GWAS, we used cases from the Osaka Acute Coronary Insufficiency Study (OACIS), which was a study that examined patients with MI at 25 collaborating hospitals in Osaka, Japan, from April 1998 to April 2006. All controls were obtained from BBJ and patients with MI, unstable angina pectoris, stable angina pectoris, cerebral infarction, or peripheral arterial disease were excluded. The baseline characteristics for cases and control in the two Japanese GWAS are shown in Supplemental Table 1. Informed consent was obtained from all participants by following the protocols approved by their institutional ethical committees before enrollment, and the Institutional Review Board (IRB) approved this study. All samples were from patients of Japanese ancestry.

**Figure 1.**
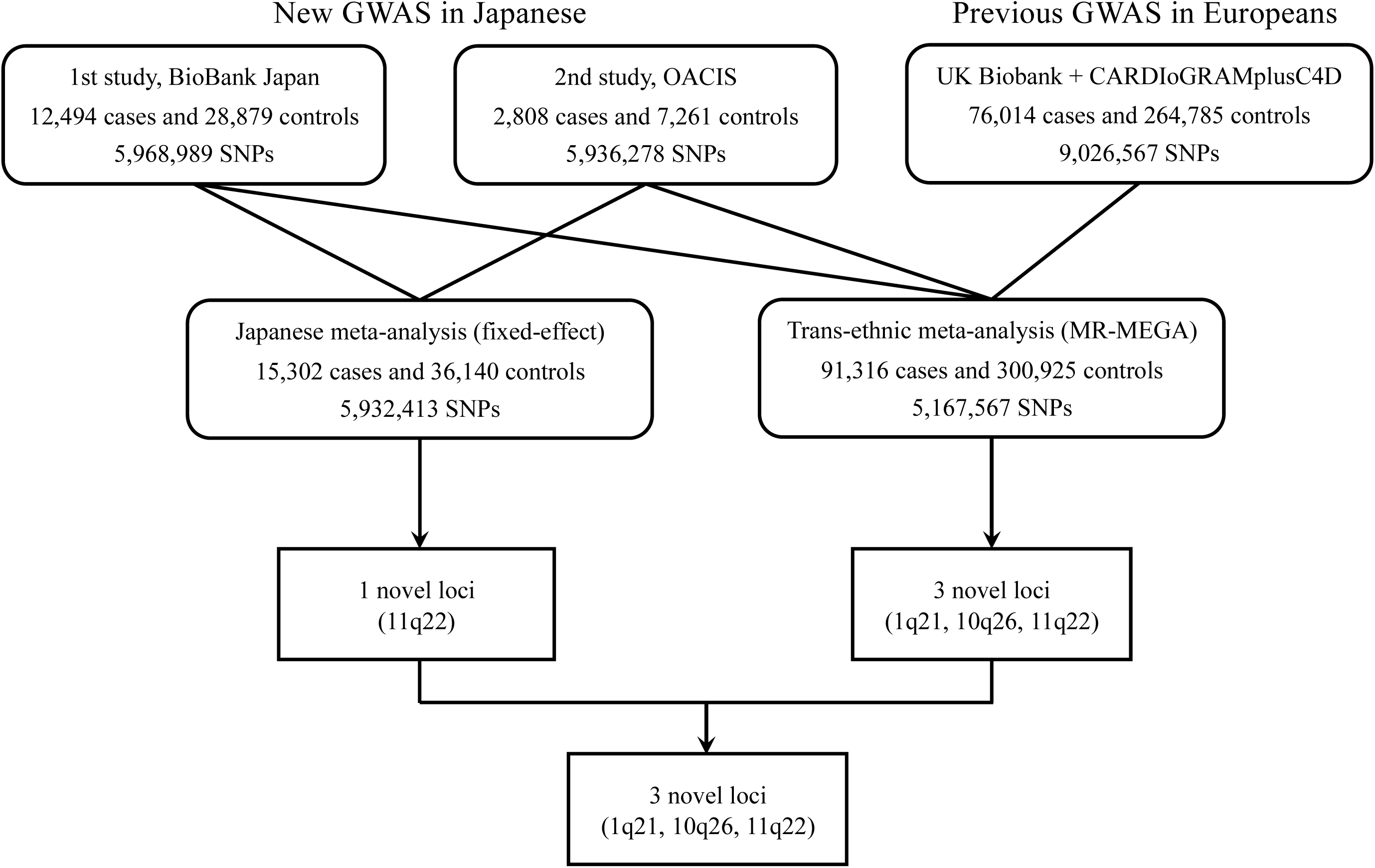
Overall design for GWAS. Two stage meta-analyses were conducted. In stage 1, Japanese meta-GWAS was performed using BioBank Japan and OACIS, which identified one new CAD loci. In stage 2, BioBank Japan, OACIS, and CARDIoGRAMplusC4D 1000 Genomes meta-analysis with UK Biobank were combined, which identified 3 novel loci. Since one locus was shared between stage 1 and stage 2, three new loci were finally identified.

### Japanese GWAS, quality control, genotyping, and imputation

We genotyped all participants using Illumina HumanOmniExpressExome BeadChip or a combination of Illumina HumanOmniExpress and HumanExome BeadChips (Illumina, Inc., San Diego, CA). SNP calling was performed by GenomeStudio software (Illumina, Inc., San Diego, CA). To perform quality control, we excluded samples with call rates < 98%. We subsequently performed analysis of identity-by-descent and identity-by-state to evaluate cryptic relatedness and removed samples if they were related to each other or duplicated in the first and second Japanese GWAS. Samples were also excluded when the genetically inferred sex did not match the reported sex. Then, we carried out principal component analysis (PCA) for the first and second Japanese GWAS to assess population stratification by using an in-house program based on the algorithm implemented by smartpca (Supplemental Figure 1). We confirmed that all cases and controls were well-matched. For single nucleotide polymorphism (SNP) quality control, we excluded variants with call rates < 99%. We also removed SNPs that deviated from the Hardy-Weinberg equilibrium with *P* < 1.0 × 10^-6^ and nonpolymorphic SNPs. Before genotype imputation, we compared the allele frequencies in our data to the reference panel based on The 1000 Genomes Project (1KG) phase I integrated release version 3 with East Asian descendants and excluded SNPs with differences > 0.16. We phased the sample haplotypes using MaCH software (ver. 1.0) and imputed missing genotypes by using Minimac software (ver.0.1.1).^12, 13^ We excluded SNPs with Rsq < 0.7 and obtained 5,968,989 and 5,936,278 SNPs in the first and second Japanese GWAS, respectively.

We performed case-control GWAS using additive logistic regression models adjusted for age, sex, dosage of effect allele (0..2), and principal components, using mach2dat (ver. 1.2.4) in the first Japanese GWAS and EPACTS (ver. 3.3.0) in the second Japanese GWAS. As for principal components, we used the first 20 components in the first Japanese GWAS and the first 10 components in the second Japanese GWAS to minimize the genomic inflation factor, lambda. We examined a quantile-quantile (Q-Q) plot and performed a single-GC correction for each GWAS. Q-Q and Manhattan plots were created using the qqman R software package (Supplemental Figure 2 and 3). Meta-analysis was performed using an inverse-variance weighted fixed-effects model implemented in METAL (2011-03-25 version). We prespecified a genome-wide significance (GWS) threshold of *P* < 5.0 × 10^-8^. To test for secondary associations near lead SNPs, we conducted conditional logistic regression analysis, where the tested SNPs were selected within 1 Mb of the lead SNPs. LocusZoom (ver. 1.3) was utilized to visualize regional information.

### Transethnic meta-analysis

We downloaded the summary statistics of the CARDIoGRAMplusC4D 1000 Genomes meta-analysis with UK Biobank data^4^ from http://www.cardiogramplusc4d.org/data-downloads/, which included 76,014 cases and 264,785 controls, mainly from European descendants, containing 9,026,567 SNPs. We used MR-MEGA software (ver. 0.1.2) for the transethnic meta-analysis.^14^ MR-MEGA provides a new approach, which demonstrated increased power over fixed- and random-effects meta-analyses, for transethnic meta-analysis across a range of scenarios of heterogeneity in allelic effects between ethnic groups. To perform the meta-analysis, the summary statistics of 3 GWAS (the first, second Japanese GWAS and CARDIoGRAMplusC4D 1000 Genomes and UK Biobank) were provided to the MR-MEGA software. Because the number of PC should be the cohort count -2, we set PC as 1 when running the software.

### Expression quantitative trait locus (eQTL), methylation quantitative trait locus (meQTL), and pleiotropy analysis

To prioritize genes associated with GWAS signals, we evaluated eQTLs obtained from the Genotype-Tissue Expression (GTEx) Portal (release v7)^15^ and meQTLs from the Biobank-Based Integrative Omics Study (BIOS).^16^ We considered the overlap between a GWAS signal and a QTL if the GWAS lead variant was in linkage disequilibrium (r^2^ ≥ 0.8 in 1KG phase I European and East Asian samples) with the lead variant of the eQTL or meQTL. Furthermore, we used the eQTL data of five subsets of immune cells (CD4^+^T cells, CD8^+^T cells, B cells, natural killer cells, and monocytes) from healthy Japanese individuals to assess gene expression in immune system-related cells thoroughly.^17^ In addition, we analyzed the GWAS results of 58 quantitative clinical traits in the Japanese population to assess clinical parameters affected by GWAS signals.^18^ The reported genes and traits were filtered using a threshold false discovery rate (FDR) of < 0.05.

### Pathway analysis

We used Meta-analysis Gene-set Enrichment of Variant Associations (MAGENTA) software (ver.2.4) to discover potential pathways associated with CAD.^19^ MAGENTA calculates *P*-values and FDRs for the gene-set enrichment analysis. The pathway databases included Gene Ontology, Kyoto Encyclopedia of Genes and Genomes (KEGG), Protein Analysis through Evolutionary Relationships (PANTHER), and Ingenuity. An FDR < 0.05 was used to evaluate the significance of associations between pathways and CAD.

### Tissue/cell-type enrichment analysis

We used Data-driven Expression-Prioritized Integration for Complex Traits (DEPICT, 2014-07-21 version) software^20^ to identify CAD-relevant tissues and cell types in the GWAS data. We prepared independent SNPs with P-values < 1 × 10^-5^ in the transethnic meta-analysis using PLINK 1.90beta software (--clump-p1 1e-5 --clump-kb 500 -clump-r2 0.05) and obtained 230 suggestive significant variants for gene prioritization. DEPICT assigned genes to associated regions if the genes resided within or were overlapping with the window in either direction with LD r^2^ > 0.5 to a given SNP. If there were no genes within the locus defined by r^2^ > 0.5, the nearest gene was selected. Using these prioritized genes, DEPICT identified enriched tissues and cell types. We set a significant threshold of FDR <0.05 for the analysis. Default settings were used in all software analysis unless otherwise noted.

## Results

### Susceptibility loci for CADs

We first conducted two Japanese GWAS comprised of 12,494 cases and 28,879 controls, and 2,808 cases and 7,261 controls, respectively. Because elevated genomic inflation factors (λ_GC_) were observed (λ_GC_ = 1.29 in the first GWAS and λ_GC_ = 1.09 in the second GWAS) despite stringent quality controls, GC corrections were applied (Supplemental Figure 2). Then, a meta-analysis of these GWAS was performed using a fixed-effects model (Figure 1). The Japanese meta-analysis included 5,932,413 variants. The λ_GC_ in the Japanese meta-GWAS was sufficiently small (λ_GC_ = 1.04) (Supplemental Figure 2). We identified a total of 18 loci that exceeded the GWS threshold of *P* < 5 × 10^-8^ in the Japanese meta-analysis (Supplemental Table 2), without evidence of heterogeneity of effects (*P_het_* > 0.10). Of these loci, 16 had been previously reported, specifically chromosome 1q41 (*MIA3*), 6p24 (*PHACTR1*), 6q23 (*TCF21*), 9p21 (*CDKN2B-AS1*), 9q33 (*DAB2IP*), 9q34 (*ABO*), 10q24 (*NT5C2*), 11q22 (*DYNC2H1-PDGFD*), 12q22 (*NR2C1*), 12q24 (*CUX2*), 13q34 (*COL4A1*), 15q25 (*ADAMTS7-MORF4L1*), 15q26 (*FES*), 19p13 (*LDLR*), 19q13 (*B9D2*) and 19q13 (*APOE*). Hence, 11q22 (*FDX1*-*ARHGAP20*) and 12q24 (*MYO1H*) were novel candidate loci. Although the lead SNP rs11066542 on chromosome 12q24 had no known loci within 1 MB, the signal completely disappeared (Supplemental Figure 4) after conditional analysis adjusted for a proxy variant of the previously reported *BRAP-ALDH2* loci (within 2 MB). Accordingly, we identified one new locus where the lead variant was rs1848599 on chromosome 11q22.

To further identify genetic loci associated with CAD, we performed a transethnic meta-analysis by MR-MEGA using our two, GC-corrected Japanese GWAS and the European data from the CARDIoGRAMplusC4D 1000 Genomes meta-analysis with UK Biobank data. In the transethnic meta-analysis, 5,167,567 variants were included. We observed 3,581 GWS variants clustering in 76 loci (Supplemental Table 3), replicating 73 previously reported loci, but finding 3 novel loci (Table 1). rs10488763, a lead variant identified in this transethnic meta-analysis and rs1848599, a GWS lead variant in the Japanese meta-analysis, were within 1 MB. We conducted conditional analyses using our two Japanese GWAS datasets and revealed that rs10488763 and rs1848599 were located within the same loci since they did not reach GWS after conditioning (Supplemental Figure 4). Collectively, 3 new loci, chromosome 1q21 (*CTSS*), 10q26 (*WDR11-FGFR2*), and 11q22 (*RDX*-*FDX1*), were discovered in our analyses.

**Table 1.**
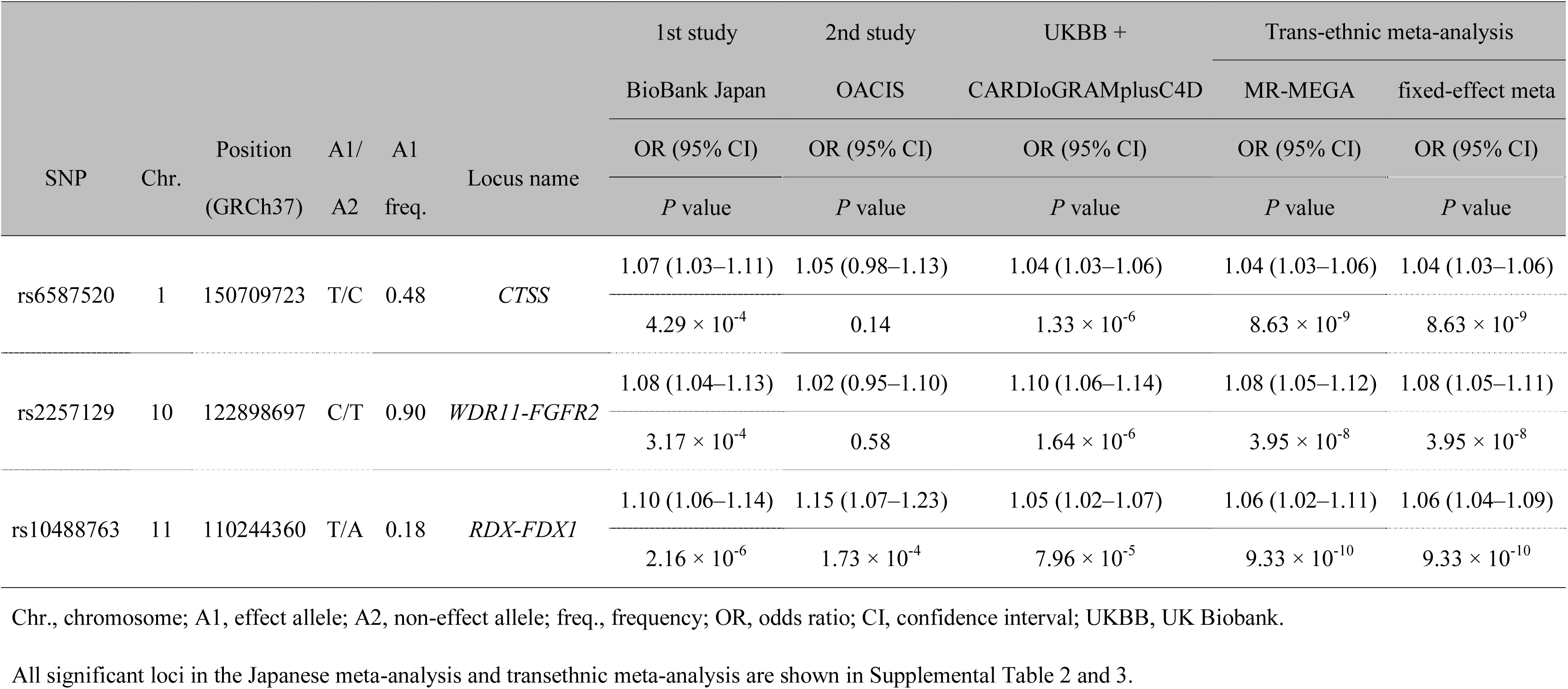
New loci identified in the transethnic meta-analysis.

### Candidate genes and biological and clinical insights of newly identified loci

We assessed the potential biology of the 3 newly identified loci by prioritizing candidate causal genes using the databases of eQTL in various cell types (GTEx) and peripheral blood cells and for DNA methylation in whole blood (meQTL:BIOS). Epigenetic annotations of these loci were explored using HaploReg. We also investigated the associations of these variants with clinical measurements using the Japanese Encyclopedia of Genetic Associations by Riken (JENGER). Additionally, we examined pleiotropy using the GWAS catalog database in order to find associations with other traits (Table 2, Supplemental Table 4).

**Table 2.**
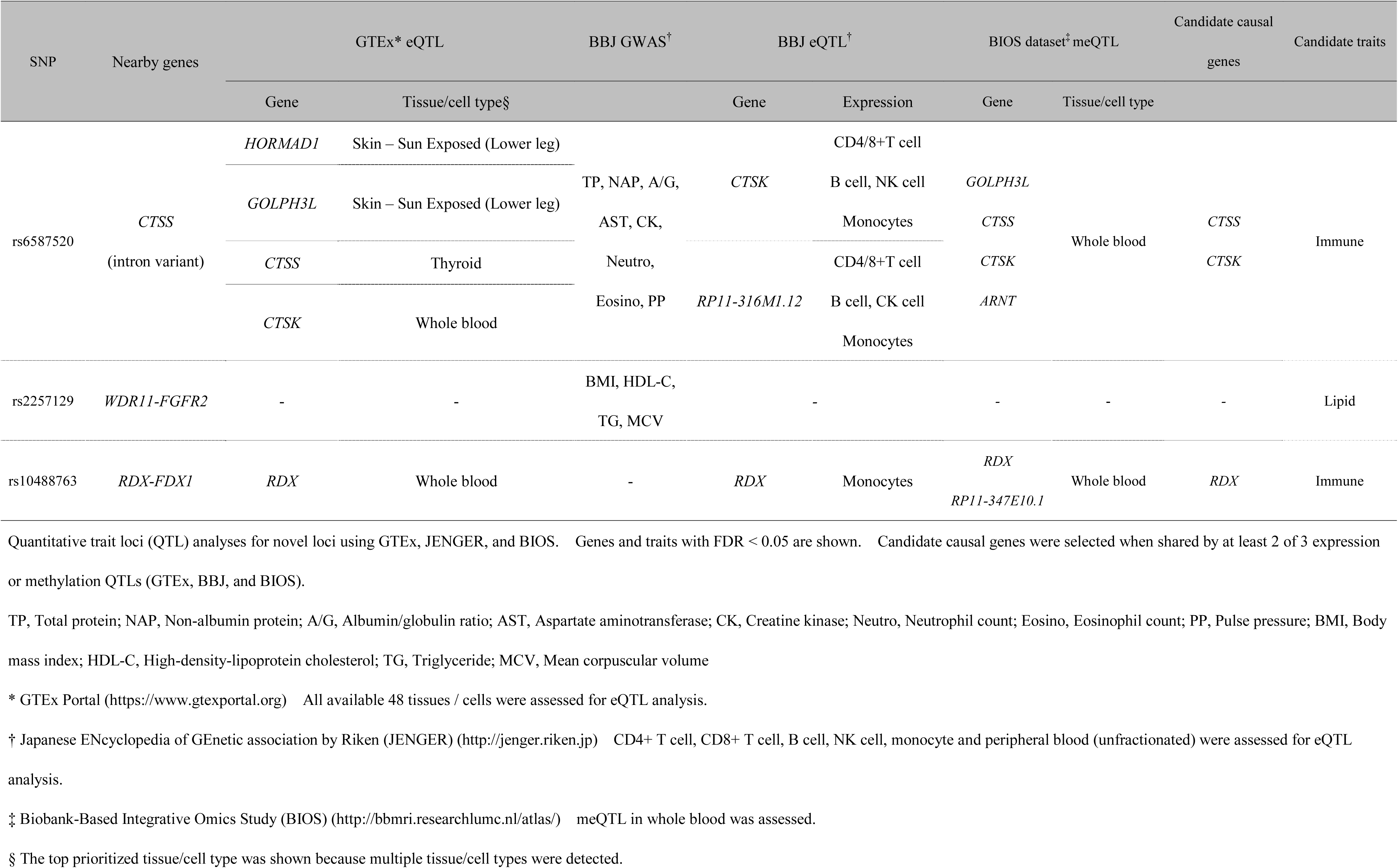
Summary of functional annotations for newly identified CAD loci.

1q21: The lead variant, rs6587520, is in an intron of *CTSS*, which encodes cathepsin S, a lysosomal cysteine proteinase that participates in the degradation of antigenic proteins to peptides for presentation on MHC class II molecules. Previous studies have reported that cathepsins play a pivotal role in inflammation, which promotes the development of atherosclerosis, MI, and hypertension.^21, 22^ We found that rs6587520 was an eQTL for *CTSS* in the left ventricle and aorta, and *CTSK* (cathepsin K) in whole blood (GTEx). Additionally, eQTL analysis of immune cells from healthy Japanese individuals showed associations of the variant with *CTSS* expression in immune cells (CD4^+^T cells, CD8^+^T cells, B cells, NK cells, and monocytes). Consistent with this finding, pleiotropy analysis based on the GWAS catalog and the 58 quantitative clinical traits in the Japanese population identified associations between this variant and several immune-cell traits, such as neutrophil and monocyte percentages of leukocytes, lymphocyte counts, and granulocyte percentage of myeloid white blood cells.

10q26: The lead variant, rs2257129, is located in an intergenic region between *WDR11* and *FGFR2*. Although the variant overlaps an H3K4me1 peak, which indicates the location of an active or primed enhancer, in several tissues including adipose and smooth muscle, we did not identify eQTLs or meQTLs associated with this variant. However, there are associations of the risk allele C with decreased high-density lipoprotein cholesterol levels and elevated triglyceride levels in plasma (Supplemental Table 5), which are traditional CAD risk factors. Apart from lipid-related traits, this risk allele has been reported to be associated with increased body mass index, which is another traditional CAD risk factor.

11q22: The lead variant, rs10488763, lies in an intergenic region between *RDX* and *FDX1*. No histone marks were found in the variant, while rs1443120 (r^2^ = 1 with the lead variant in Europeans and East Asians) overlaps promoter histone marks in fat and blood cells. There are associations of the risk allele T with increased expression of *RDX* in whole blood (GTEx) and monocytes (JENGER). Furthermore, meQTL analyses revealed associations with *RDX* in whole blood (BIOS). *RDX* encodes radixin, which is a cytoskeletal protein that plays an important role in connecting actin to the plasma membrane. A previous report suggested that radixin controls vascular smooth muscle cell migration, which is involved in pathophysiological processes such as angiogenesis and atherosclerosis.^23^ However, clinical measurements in the Japanese population and pleiotropy analyses showed no associated traits or diseases.

### Pathway analysis and CAD-related tissues and cell types

Using the result of the transethnic meta-analysis, we investigated biological pathways associated with CAD using MAGENTA software. We found 25 significant biological pathways (FDR < 0.05, Supplemental Table 6). Lipid-related pathways were most frequent (13/25), which was followed by transport (4/25), signaling (3/25) pathways. These results suggest that the development of CAD could involve a variety of biological processes, including lipid metabolism/transport/homeostasis, inflammation, cell signaling/cycle/proliferation and regulation of blood vessel.

Next, we performed tissue/cell-type enrichment analysis using DEPICT software (Table 3). The most significant associations were observed in arteries (*P* = 3.05 × 10^-6^). Moreover, we found a significant enrichment in adrenal glands (*P* = 1.31 × 10^-5^), which are endocrine glands and play an important role in the regulation of metabolism, immune function and blood pressure. We also found enrichment in fat tissues, confirming the association of lipid metabolism with CAD.

**Table 3.** Tissue/cell type enrichment analysis by DEPICT.

| Name | MeSH first level term | MeSH second level term | *P* value | FDR < 5% |
| --- | --- | --- | --- | --- |
| Arteries | Cardiovascular System | Blood Vessels | $3.05 \times 10^{-6}$ | Yes |
| Adrenal Glands | Endocrine System | Endocrine Glands | $1.31 \times 10^{-5}$ | Yes |
| Adipose Tissue | Tissues | Connective Tissue | $1.78 \times 10^{-5}$ | Yes |
| Adrenal Cortex | Endocrine System | Endocrine Glands | $2.85 \times 10^{-5}$ | Yes |
| Subcutaneous Fat | Tissues | Connective Tissue | $3.13 \times 10^{-5}$ | Yes |
| Adipose Tissue, White | Tissues | Connective Tissue | $3.13 \times 10^{-5}$ | Yes |
| Subcutaneous Fat, Abdominal | Tissues | Connective Tissue | $3.84 \times 10^{-5}$ | Yes |
| Abdominal Fat | Tissues | Connective Tissue | $3.84 \times 10^{-5}$ | Yes |
| Adipocytes | Cells | Connective Tissue Cells | $1.88 \times 10^{-4}$ | Yes |
| Stomach | Digestive System | Gastrointestinal Tract | $2.27 \times 10^{-4}$ | Yes |

DEPICT, Data-driven Expression-Prioritized Integration for Complex Traits.
MeSH, Medical Subjects Heading. FDR, false discovery rate.
Top 10 tissues/cell types are shown.

### Susceptibility differences for CAD between Japanese and Europeans

We investigated the differences in genetic susceptibility to MI between Japanese and Europeans. First, we compared all variants shared by both our Japanese meta-analysis and the CARDIoGRAMplusC4D 1000 Genomes MI subphenotype analysis, where both cases were comprised of MI patients (Supplemental Figure 5). While we found some common signals between those two studies, population-specific signals were also observed. Next, using 76 lead variants identified in our transethnic meta-analysis, we compared the odds ratios of those variants (Figure 3). The odds ratios were very similar in most of the variants. We found a moderate correlation of odds ratios between Japanese and Europeans (Spearman’s ρ= 0.56, *P* = 1.21 × 10^-7^). However, the odds ratios for rs11124924, rs1250229, rs3130342, rs2107595, rs643434, rs1558803, rs7139170, and rs62010554 were markedly different between the two cohorts as their 95% confidence intervals did not reach the line of equality. This result suggests that the *ABO* locus, which harbors rs643434, is preferentially linked to MI in Europeans, while the other 7 loci, *APOB*, *FN1*, *ATF6B, HDAC9*, *UBE3B, RPH3A,* and *ADAMTS7*, are preferentially linked to MI in Japanese.

**Figure 2.**
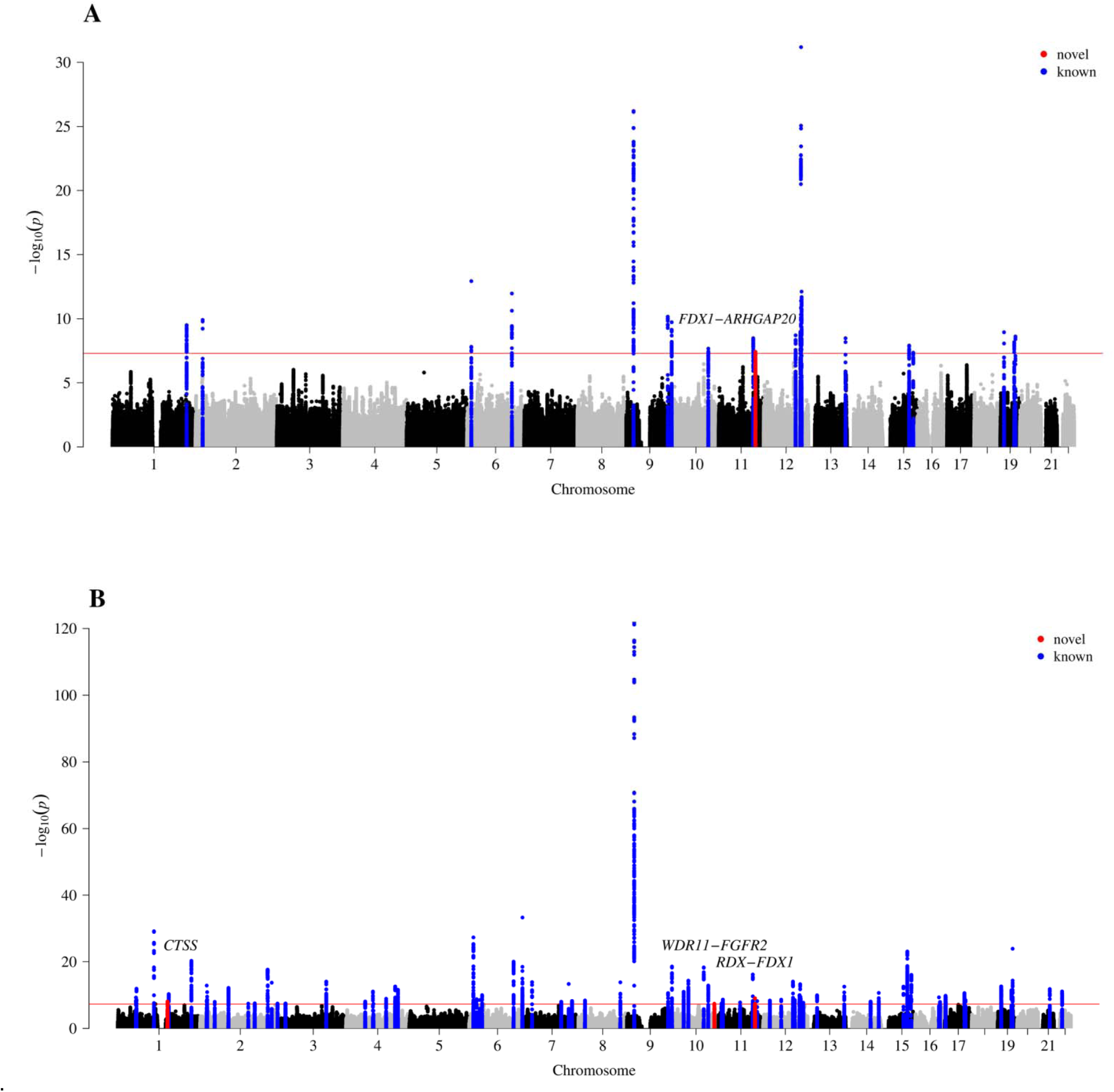
Manhattan plots in the Japanese meta-analysis and transethnic meta-analysis. Manhattan plot for **A**, Japanese meta-GWAS (15,302 CAD cases and 36,140 controls), **B**, Transethnic meta-analysis (91,316 CAD cases and 300,925 controls). The *x* axis denotes chromosomal location and *y* axis denotes -log_10_ *P* value for each SNP. The horizontal red line shows a threshold of GWS (*P* = 5.0 × 10^-8^). Red and blue dots represent novel and known GWS loci (lead variants ± 250 kb), respectively.

**Figure 3.**
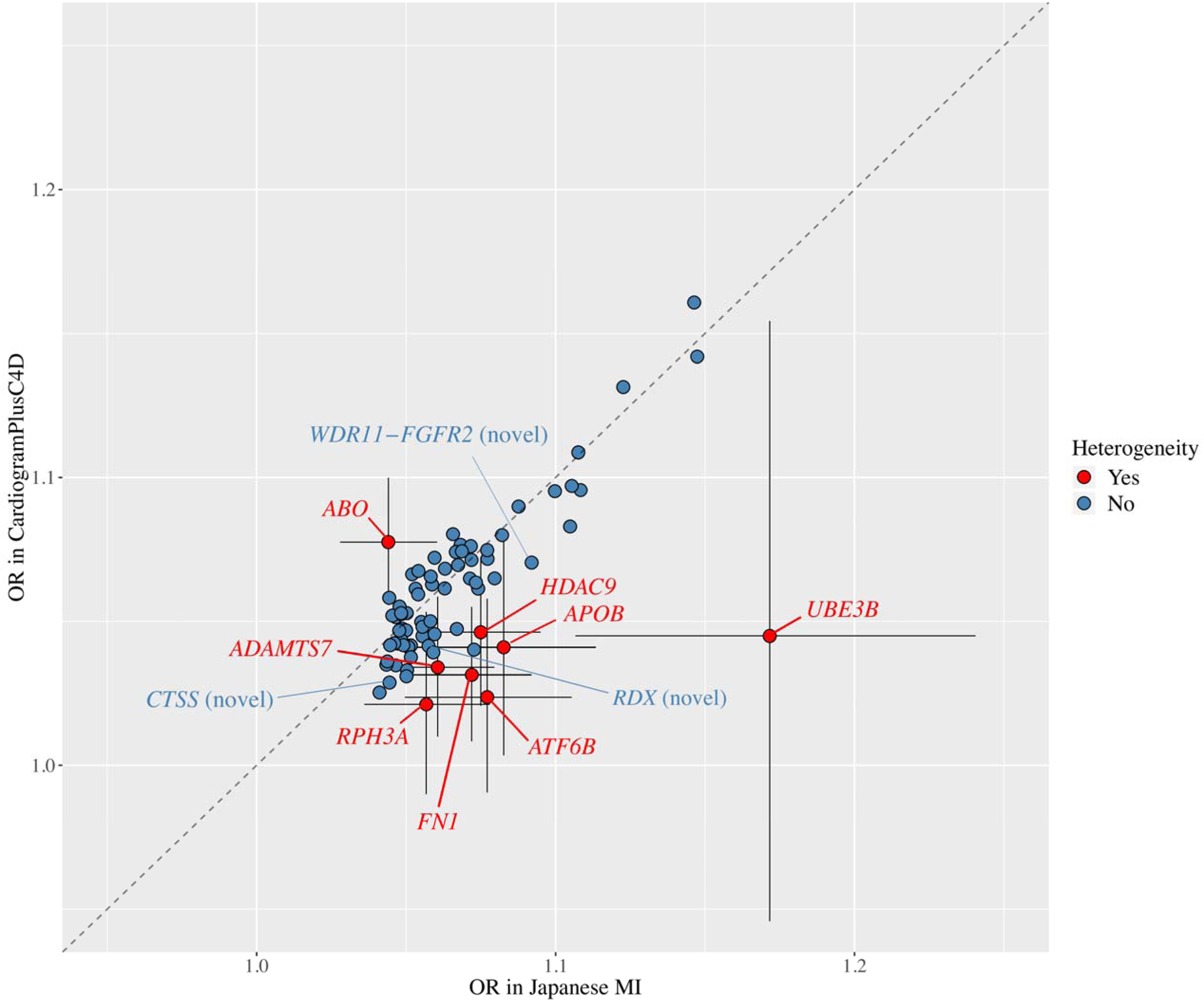
Odds ratios of lead variants for MI between Japanese and Europeans. Scatter plot of odds ratios (ORs) between Japanese (*x* axis) and CardiogramPlusC4D (*y* axis) for MI using 76 lead variants in the transethnic meta-analysis. Variants whose 95% confidence interval did not reach the line of equality (*y* = *x*) are shown in red. Error bars represent the 95% confidence interval for each variant.

Next, we compared the risk allele frequency (RAF) of those variants in 1KG phase3 between East Asians, to which Japanese belong, and Europeans, in order to observe the distributions in a population cohort (Figure4, Supplemental Table 7). We first confirmed strong correlations of RAFs between our Japanese meta-analysis and 1KG phase3 East Asians (Spearman’s ρ= 0.98, *P* = 1.50 × 10^-52^), and the CARDIoGRAMplusC4D 1000 Genomes meta-analysis with UK Biobank data and 1KG phase 3 Europeans (Spearman’s ρ= 0.99, *P* = 2.71 × 10^-59^), respectively. Then, we found that there was no significant imbalance in the number between East Asian-frequent variants and European-frequent variants (*P* for sign test = 0.91). When we assessed the 3 lead variants for newly identified loci, we found just 1 variant was more frequent for East Asians (rs10488763: 37% in East Asians vs. 11% in Europeans), while the others were more frequent for Europeans (rs6587520: 46% in East Asians vs 48% in Europeans, rs2257129: 61% in East Asians vs 97% in Europeans). In contrast to the RAFs, rs10488763 did not reach a suggestive significance level (*P* < 1 x 10^-5^) in the CARDIoGRAMplusC4D 1000 Genomes meta-analysis with UK Biobank, while the others reached. Taken together, both more frequent risk allele frequency in East Asians and increased sample size may help to detect these new signal.

**Figure 4.**
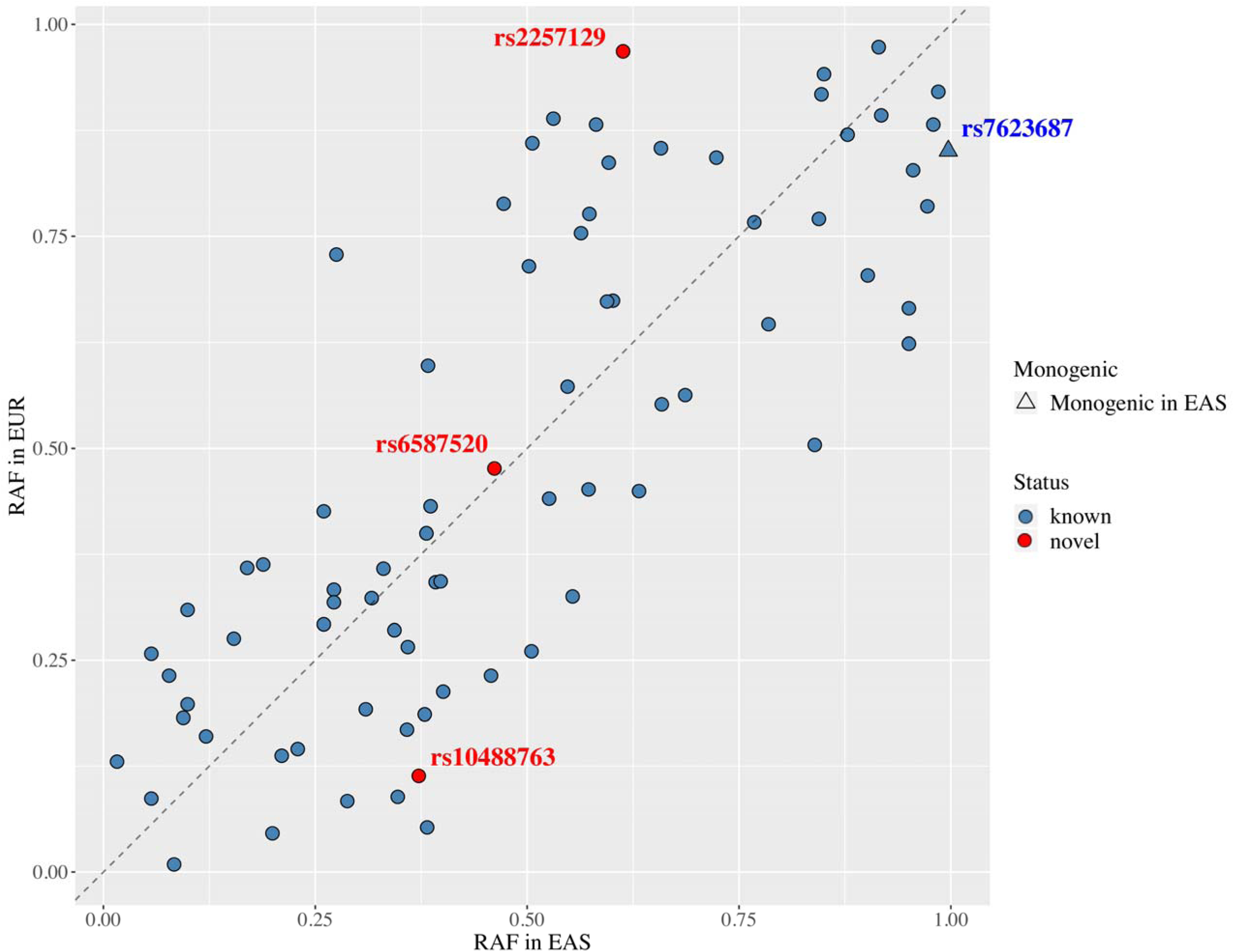
Distribution of RAFs of 77 lead variants between East Asians and Europeans. Scatter plot of risk allele frequencies (RAFs) between East Asians (x axis) and Europeans (y axis) for MI using 76 lead variants in our transethnic meta-analysis and 1 lead variant (rs7623687) found only in the CARDIoGRAMplusC4D 1000 Genomes-based GWAS. Although rs7623687 showed a genome-wide significant association with MI in the CARDIoGRAMplusC4D 1000 Genomes-based GWAS, we did not use this variant because it was monomorphic in East Asians. RAFs were based on 1000 Genomes Project Phase 3. Novel loci are shown in red. Circles denote 76 lead variants of genome-wide significant loci in our transethnic meta-analysis, while a triangle denotes a monogenic locus in East Asians. EAS, East Asians; EUR, Europeans.

Finally, we divided 230 suggestive significant variants used in the tissue/cell-type enrichment analysis into two groups, the East Asian-frequent group and the European-frequent group, according to the RAF in 1KG phase3 (Supplemental Table 8). We then removed variants with the absolute value of [RAF in East Asian - RAF in European] <0.05 because variants with small RAF difference do not characterize the population-specificity. As a result, the East Asian-frequent group had 93 variants, while the European-frequent group had 92 variants; Using these variants, we again performed tissues/cell type enrichment analysis and found different types of enrichment (Figure 5, Supplemental Table 9). The East Asian-frequent group showed significant enrichment in adrenal glands and adrenal cortex, while there was a significant enrichment in arteries and fat tissues in the European-frequent group. Variations in RAF can contribute to differences in the prevalence of diseases among populations.^24^ Thus, these findings suggest that there are differences in CAD susceptibility between Japanese and Europeans.

**Figure 5.**
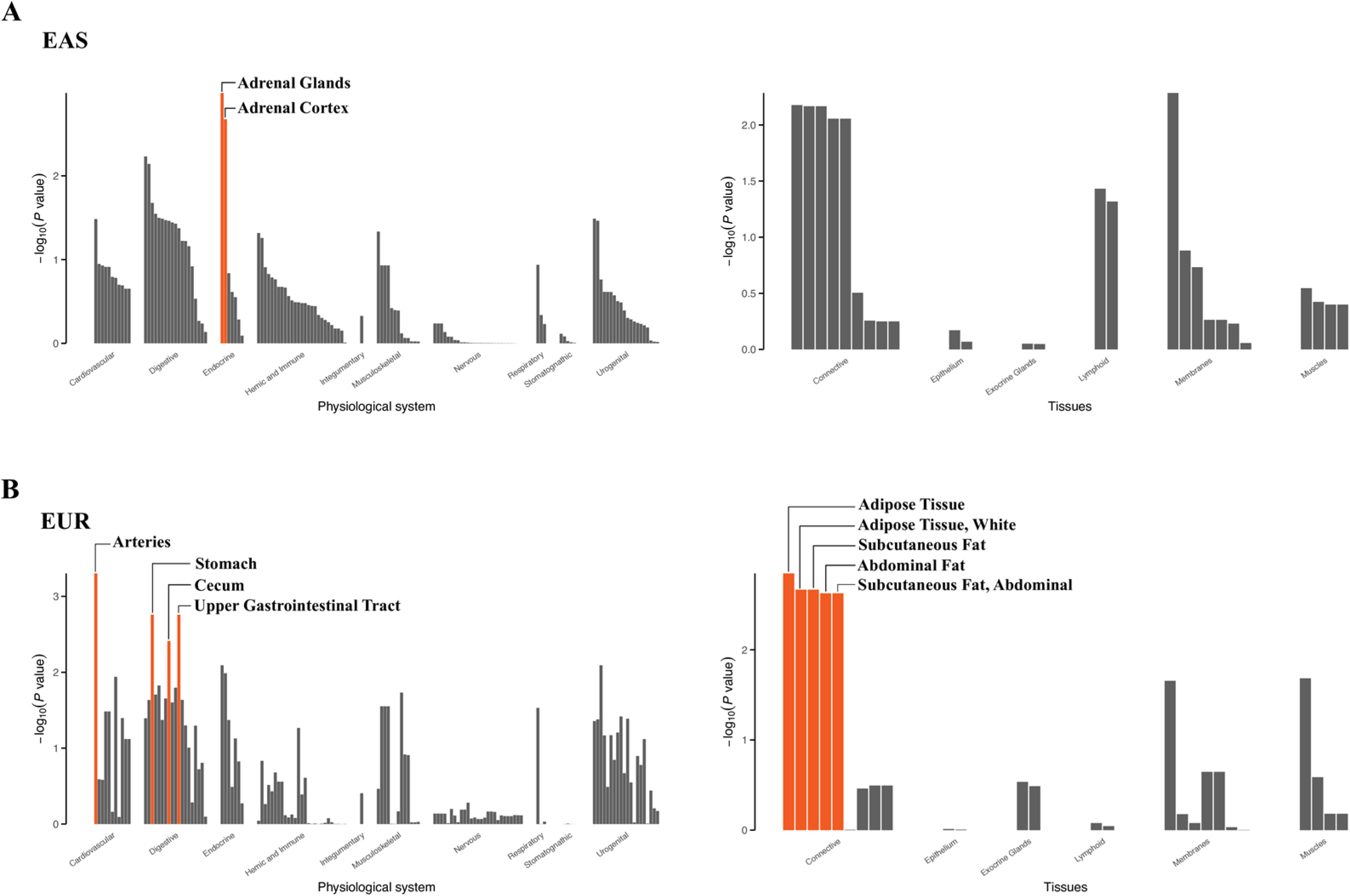
Population-specific tissue/cell type enrichment analysis by DEPICT. Results of tissue/cell type enrichment analysis for A. the East Asian-frequent group and B. the European-frequent group. The *x* axis shows physiological systems and tissues grouped by Medical Subject Heading (MeSH) terms. The *y* axis shows -log_10_ *P* value for each tissue/cell type. The orange bars represent a significantly enriche tissue/cell type (FDR<0.05).

## Discussion

We conducted the largest Japanese GWAS for CAD to date, which was comprised of 15,302 cases and 36,140 controls. Additionally, we performed a transethnic meta-analysis with the CARDIoGRAMplusC4D 1000 Genomes meta-analysis with UK Biobank data, including 88,192 cases and 162,544 controls. We finally validated 73 previously reported loci and identified 3 new loci for CAD at the GWS level. Using the lead variants in the identified loci, we compared odds ratios for MI and RAFs to elucidate the genetic differences between Japanese and Europeans. Akiyama et al. reported that the lead variant rs2257129 in the new loci *WDR11-FGFR2* is associated with body mass index,^25^ which is a clinical measure of obesity. Because obesity is a conventional risk factor for CAD, it seems natural to suppose that the locus linked to obesity is associated with the development of CAD. Additionally, clinical QTL analyses using the JENGER database suggested that rs2257129 is associated with triglyceride and high-density lipoprotein cholesterol levels in the blood, which are also known as clinical laboratory values associated with CAD. Actually, a recent GWAS for CAD reported that this variant reached a threshold for suggestive significance (5e-7).^6^ However, our analyses showed no significant eQTL or meQTL genes for rs2257129, which could have explained a molecular mechanism between the genetic variation and phenotype. Further research is necessary to improve our understanding of how rs2257129 affects body mass index, triglycerides, high-density lipoprotein cholesterol, and the development of CAD.

The findings from eQTL and meQTL analyses indicated that rs6587520, the lead variant in the *CTSS* locus, modulates the expression of cathepsin genes (i.e., *CTSS* and *CTSK*) in immune cells. Cathepsins are lysosome-restricted proteases, which have been reported to be related to many cardiovascular diseases, such as atherosclerosis, CAD, cardiac hypertrophy, cardiomyopathy, and hypertension. Cathepsin S (CTSS) and cathepsin K (CTSK) were the first cathepsins found in endothelial cells, macrophages, and smooth muscle cells from human atherosclerotic lesions.^26^ In *ApoE*^-/-^ mice, an atherosclerotic murine model, knockdown of *Ctss* produced smaller plaque sizes and fewer acute plaque ruptures.^27^ In addition, active cathepsin S was localized preferentially to macrophages and was increased in the areas of disrupted elastin fibers.^26^ Moreover, in the *Ldlr*^-/-^ atherosclerotic mouse model, knockdown of *Ctss* reduced the percentage of plaque area composed of monocytes/macrophages.^28^ These findings suggest that cathepsin S and K influence the progression of CAD via the development of atherosclerosis through the immune system.

eQTL analysis also suggested that rs10488763, the lead variant in the *FDX1* locus, alters the expression of *RDX* in monocytes. Radixin, the protein *RDX* encodes, is a member of the ERM (ezrin/radixin/moesin) family, which plays a pivotal role in interactions between the plasma membrane and cytoskeleton. ERM family members are involved in cell migration and adhesion. Previous studies have shown that inflammatory stimuli, such as angiotensin II and interleukin-1ß, can increase Rho-kinase expression in cultured human coronary vascular smooth muscle cells. Then, the elevation of Rho-kinase expression leads to increased phosphorylation of EMR family members, resulting in the progression of atherosclerosis.^29^ Consequently, radixin, one of the EMR family members, may be involved in the pathogenesis of atherosclerosis via the immune system. Given the results of the QTL analyses, two of the 3 novel loci, *CTSS* and *FDX1,* are suggested to be involved in the immune system, which plays an important role in the development of atherosclerosis. Hence, our new loci highlight the involvement of the immune-atherosclerosis axis in the development of CAD.

Pathway analysis showed that lipid metabolism pathway was the most relevant for CAD. Furthermore, tissue/cell-type enrichment analysis confirmed the involvement of fat tissues such as subcutaneous fat and abdominal fat in CAD. In addition, our analysis suggested the involvement of adrenal glands in the disease biology. The adrenal glands play a multi-functional role in the endocrine system and produce a variety of hormones such as catecholamines and corticosteroids. Catecholamines regulate the heart rate and blood pressure, while corticosteroids influence stress response, immune response, regulation of inflammation, and carbohydrate metabolism. These findings suggest an endocrine system regulated by the adrenal glands plays an important role in the development of CAD, as well as lipid metabolism.

The majority of previous GWAS for CAD were conducted in European populations,^3–5^ and we found a relatively small number of GWAS performed in non-European populations.^1, 2^ A previous study suggested that the directions of effect in CAD-related loci were generally concordant between Europeans and non-Europeans,^30^ and our study also demonstrated that the directions and sizes of the effect of the lead variants detected in our transethnic meta-analysis were almost the same between Japanese and Europeans. However, 7 out of the 76 variants had a greater effect and one had a lesser effect in the Japanese population. Aside from the effect sizes, we found different distributions in the RAFs of those variants between Japanese and Europeans. Especially, the RAFs in 1 new locus, rs10488763, was higher in East Asians than in Europeans, which may have resulted in exceeding a GWS threshold in our transethnic meta-GWAS encompassing more East Asians than in previous studies. Since there are differences in genetic architecture across populations, it is crucial to perform GWAS for CAD in other populations.

In the tissue/cell-type enrichment analysis for each population, we identified significant enrichment of adrenal glands in East Asians, while we found an enrichment of arteries and fat tissues for Europeans. This difference could explain different susceptibilities to CAD between East Asians and Europeans from a biological perspective. With respect to fat tissues—which is significantly associated with atherosclerosis—East Asians including Japanese have the lowest amount of subcutaneous fat compared to other populations, suggesting that lipid spillover can easily occurr.^31^ Consistent with this, it was reported that Japanese Americans had greater levels of atherosclerosis compared to European Americans,^32^ despite the lower rate of CAD in Japan. This report implies that genetically, Japanese are not less prone to atherosclerosis. As such, these findings could explain a part of the difference in the genetic basis of CAD between East Asians and Europeans.

Our study has some limitations. First, we adopted the CARDIoGRAMplusC4D 1000 Genomes meta-analysis with UK Biobank data for the CAD study in European populations. While most of these samples (∼91%) are of European descent, a small portion is of East Asian descent. This could lead to underestimation of the differences in the effect size in CAD loci between Japanese and Europeans. However, we were successful in identifying several differences despite the conservative settings. Second, the sample size in our Japanese study was modest compared to that of recent CAD GWAS, which resulted in a smaller number of new loci. Because population specificity is important in genetic analysis, greater sample sizes will be required in the future for further understanding.

In summary, we have reported 3 new loci for CAD. Our results have provided a well-powered CAD GWAS for East Asians, which could be useful to implement precision medicine (e.g. developing an East-Asian specific polygenic risk score for CAD). Moreover, our transethnic analyses showed both shared and unique genetic architectures between the Japanese and European populations. Since there are different genetic architectures across populations, further non-European analyses will be needed to elucidate the variety in genetic architecture of CAD.

## Acknowledgements

We are grateful to the members of BioBank Japan and the Rotary Club of Osaka-Midosuji District for supporting our study. This work was conducted as a part of the BioBank Japan Project.

## Funding Sources

The BioBank Japan Project was supported by the Ministry of Education, Culture, Sports, Science and Technology, Japan. The JPHC Study was supported by National Cancer Center Research and Development Fund since 2011 and a grant-in-aid for Cancer Research from the Ministry of Health, Labour and Welfare of Japan from 1989 to 2010. This research was funded by the GRIFIN project of Japan Agency for Medical Research and Development.

## Disclosures

None

