## Supplemental Materials for "Transethnic meta-analysis of genome-wide association studies identifies three new loci and characterizes population-specific differences for coronary artery disease"

### SUPPLEMETAL MATERIAL

Supplemental Table 1. Baseline Characteristics for cases and controls in the Japanese GWASs

| | 1st Japanese GWAS | | 2nd Japanese GWAS | |
|---|---|---|---|---|
| | Cases (N=12,494) | Controls (N=28,879) | Cases (N=2,808) | Controls (N=7,261) |
| Male | 80.7% | 39.4% | 78.4% | 57.8% |
| Age (mean±s.d.) | 66.9±10.5 | NA | 64.8±11.5 | 63.0±9.8 |
| BMI | 23.9±3.4 | NA | 23.9±3.5 | 22.4±3.7 |
| Medical history | | | | |
| Dyslipidemia | 52.1% | NA | 45.6% | 12.5% |
| Hypertension | 53.4% | NA | 60.6% | 50.0% |
| Type 2 diabetes mellitus | 34.7% | NA | 33.2% | 10.1% |

NA, not available.

**Supplemental Table 2. Significant loci in the Japanese meta-analysis**

| SNP | Chr. | Position (GRCh37) | A1/ A2 | A1 freq | Locus name | Status | 1st study, BioBank Japan OR (95% CI) | 1st study, BioBank Japan *P* value | 2nd study, OACIS OR (95% CI) | 2nd study, OACIS *P* value | Japanese meta-analysis OR (95% CI) | Japanese meta-analysis *P* value |
|---|---|---|---|---|---|---|---|---|---|---|---|---|
| rs67180937 | 1 | 222823743 | G/T | 0.54 | *MIA3* | Known | 1.14 (1.10—1.18) | $5.39 \times 10^{-11}$ | 1.09 (1.01—1.17) | 0.019 | 1.13 (1.09—1.17) | $3.18 \times 10^{-10}$ |
| rs2876301 | 6 | 12907411 | T/C | 0.81 | *PHACTR1* | Known | 1.17 (1.12—1.23) | $5.22 \times 10^{-9}$ | 1.12 (1.01—1.23) | 0.026 | 1.15 (1.09—1.21) | $1.72 \times 10^{-8}$ |
| rs2327429 | 6 | 134209837 | T/C | 0.48 | *TCF21* | Known | 1.13 (1.09—1.17) | $1.30 \times 10^{-10}$ | 1.17 (1.09—1.26) | $9.34 \times 10^{-6}$ | 1.14 (1.10—1.18) | $1.09 \times 10^{-12}$ |
| rs10757274 | 9 | 22096055 | G/A | 0.48 | *CDKN2B-AS1* | Known | 1.23 (1.19—1.28) | $1.59 \times 10^{-26}$ | 1.21 (1.13—1.30) | $2.48 \times 10^{-7}$ | 1.22 (1.18—1.27) | $6.33 \times 10^{-27}$ |
| rs10985348 | 9 | 124419722 | C/G | 0.24 | *DAB2IP* | Known | 1.16 (1.11—1.22) | $5.07 \times 10^{-11}$ | 1.13 (1.04—1.22) | $3.67 \times 10^{-3}$ | 1.16 (1.11—1.21) | $7.01 \times 10^{-11}$ |
| rs9411378 | 9 | 136145425 | A/C | 0.26 | *ABO* | Known | 1.13 (1.08—1.18) | $3.28 \times 10^{-7}$ | 1.22 (1.13—1.32) | $7.13 \times 10^{-7}$ | 1.15 (1.10—1.20) | $1.89 \times 10^{-10}$ |
| rs11191559 | 10 | 104867686 | C/T | 0.73 | *NT5C2* | Known | 1.15 (1.10—1.20) | $7.81 \times 10^{-10}$ | 1.07 (0.99—1.16) | $8.40 \times 10^{-2}$ | 1.13 (1.08—1.18) | $2.17 \times 10^{-8}$ |
| rs11226017 | 11 | 103664409 | C/T | 0.63 | *DYNC2H1-PDGFD* | Known | 1.14 (1.10—1.18) | $4.77 \times 10^{-11}$ | 1.07 (0.99—1.15) | $8.90 \times 10^{-2}$ | 1.13 (1.08—1.17) | $3.32 \times 10^{-9}$ |
| rs1848599 | 11 | 110358465 | C/T | 0.78 | *FDX1-ARHGAP20* | Novel | 1.12 (1.06—1.17) | $2.60 \times 10^{-6}$ | 1.19 (1.09—1.29) | $9.24 \times 10^{-5}$ | 1.14 (1.09—1.19) | $3.87 \times 10^{-8}$ |
| rs933899 | 12 | 95446288 | A/G | 0.77 | *NR2C1* | Known | 1.15 (1.10—1.20) | $1.27 \times 10^{-9}$ | 1.13 (1.03—1.23) | $8.44 \times 10^{-3}$ | 1.15 (1.10—1.20) | $1.97 \times 10^{-9}$ |
| rs79105258 | 12 | 111718231 | A/C | 0.26 | *CUX2* | Known | 1.34 (1.28—1.40) | $1.26 \times 10^{-33}$ | 1.25 (1.15—1.35) | $5.28 \times 10^{-8}$ | 1.31 (1.25—1.37) | $6.55 \times 10^{-32}$ |
| rs4773140 | 13 | 110954237 | G/A | 0.70 | *COL4A1* | Known | 1.14 (1.09—1.19) | $1.84 \times 10^{-9}$ | 1.11 (1.02—1.19) | $8.62 \times 10^{-3}$ | 1.14 (1.09—1.19) | $3.33 \times 10^{-9}$ |
| rs28455815 | 15 | 79114453 | T/C | 0.54 | *ADAMTS7-MORF4L1* | Known | 1.12 (1.07—1.16) | $5.63 \times 10^{-8}$ | 1.12 (1.04—1.20) | $1.57 \times 10^{-3}$ | 1.12 (1.08—1.16) | $1.24 \times 10^{-8}$ |
| rs4533264 | 15 | 91431524 | T/C | 0.20 | *FES* | Known | 1.15 (1.10—1.21) | $1.41 \times 10^{-8}$ | 1.10 (1.01—1.20) | $2.40 \times 10^{-2}$ | 1.14 (1.09—1.19) | $4.50 \times 10^{-8}$ |
| rs2738464 | 19 | 11242307 | C/G | 0.68 | *LDLR* | Known | 1.14 (1.09—1.19) | $7.36 \times 10^{-10}$ | 1.12 (1.03—1.20) | $5.01 \times 10^{-3}$ | 1.13 (1.08—1.17) | $1.15 \times 10^{-9}$ |
| rs4803458 | 19 | 41865293 | A/G | 0.53 | *B9D2* | Known | 1.11 (1.06—1.15) | $2.94 \times 10^{-7}$ | 1.15 (1.07—1.23) | $1.67 \times 10^{-4}$ | 1.12 (1.08—1.16) | $6.66 \times 10^{-9}$ |
| rs769446 | 19 | 45408628 | T/C | 0.95 | *APOE* | Known | 1.34 (1.21—1.47) | $8.71 \times 10^{-9}$ | 1.34 (1.12—1. 59) | $1.36 \times 10^{-3}$ | 1.34 (1.22—1.47) | $2.44 \times 10^{-9}$ |

Chr., chromosome; A1, effect allele; A2, non-effect allele; freq., frequency; OR, odds ratio; CI, confidence interval.

**Supplemental Table 3. Significant loci in the transethnic meta-analysis**

| | | | | | | | 1st study, BioBank Japan | | 2nd study, OACIS | | UKBB + CARDIoGRAMplusC4D | | Transethnic meta-analysis | | | |
|---|---|---|---|---|---|---|---|---|---|---|---|---|---|---|---|---|
| | | | | | | | | | | | | | MR-MEGA | | fixed-effect meta | |
| SNP | Chr. | Position (GRCh37) | A1/ A2 | A1 freq | Locus name | Status | OR (95% CI) | *P* value | OR (95% CI) | *P* value | OR (95% CI) | *P* value | OR (95% CI) | *P* value | OR (95% CI) | *P* value |
| rs11206510 | 1 | 55496039 | T/C | 0.85 | *BSND -PCSK9* | Known | 1.14 (1.04—1.24) | $3.52 \times 10^{-3}$ | 0.96 (0.82—1.13) | 0.65 | 1.07 (1.04—1.09) | $3.68 \times 10^{-8}$ | 1.07 (1.04—1.10) | $1.01 \times 10^{-8}$ | 1.07 (1.04—1.09) | $1.01 \times 10^{-8}$ |
| rs56348932 | 1 | 56988551 | A/C | 0.92 | *PPAP2B* | Known | 1.11 (0.94—1.29) | 0.20 | 1.28 (0.96—1.72) | $9.30 \times 10^{-2}$ | 1.11 (1.07—1.14) | $4.07 \times 10^{-12}$ | 1.11 (1.08—1.13) | $2.31 \times 10^{-12}$ | 1.11 (1.07—1.14) | $2.31 \times 10^{-12}$ |
| rs4970837 | 1 | 109822008 | T/G | 0.67 | *PSRC1* | Known | 1.20 (1.10—1.30) | $8.24 \times 10^{-6}$ | 1.14 (0.99—1.31) | $6.40 \times 10^{-2}$ | 1.07 (1.05—1.09) | $1.19 \times 10^{-13}$ | 1.07 (1.04—1.11) | $7.69 \times 10^{-16}$ | 1.07 (1.06—1.09) | $7.69 \times 10^{-16}$ |
| rs6587520 | 1 | 150709723 | T/C | 0.48 | *CTSS* | Novel | 1.07 (1.03—1.11) | $4.29 \times 10^{-4}$ | 1.05 (0.98—1.13) | 0.14 | 1.04 (1.03—1.06) | $1.33 \times 10^{-6}$ | 1.04 (1.03—1.06) | $8.63 \times 10^{-9}$ | 1.04 (1.03—1.06) | $8.63 \times 10^{-9}$ |
| rs11810571 | 1 | 151762308 | G/C | 0.77 | *TDRKH* | Known | 1.05 (1.01—1.09) | $9.63 \times 10^{-3}$ | 0.96 (0.89—1.03) | 0.25 | 1.06 (1.04—1.08) | $2.21 \times 10^{-8}$ | 1.05 (1.02—1.09) | $4.41 \times 10^{-8}$ | 1.05 (1.03—1.07) | $4.41 \times 10^{-8}$ |
| rs12129500 | 1 | 154423764 | T/C | 0.46 | *IL6R* | Known | 1.06 (1.02—1.11) | $1.90 \times 10^{-3}$ | 0.97 (0.90—1.04) | 0.38 | 1.05 (1.03—1.06) | $8.52 \times 10^{-9}$ | 1.05 (1.02—1.07) | $2.84 \times 10^{-9}$ | 1.05 (1.03—1.06) | $2.84 \times 10^{-9}$ |
| rs3008629 | 1 | 222835930 | T/C | 0.80 | *MIA3* | Known | 1.12 (1.07—1.16) | $5.68 \times 10^{-8}$ | 1.07 (0.99—1.15) | $6.80 \times 10^{-2}$ | 1.06 (1.04—1.08) | $1.94 \times 10^{-7}$ | 1.07 (1.04—1.10) | $5.81 \times 10^{-12}$ | 1.07 (1.05—1.09) | $5.81 \times 10^{-12}$ |
| rs16986953 | 2 | 19942473 | A/G | 0.13 | *OSR1 -TTC32* | Known | 1.09 (1.05—1.14) | $6.59 \times 10^{-5}$ | 1.09 (1.02—1.17) | $1.60 \times 10^{-2}$ | 1.12 (1.08—1.15) | $4.77 \times 10^{-10}$ | 1.10 (1.09—1.12) | $1.39 \times 10^{-13}$ | 1.10 (1.08—1.13) | $1.39 \times 10^{-13}$ |
| rs11124924 | 2 | 21206275 | G/C | 0.92 | *LDAH -APOB* | Known | 1.16 (1.09—1.23) | $6.83 \times 10^{-7}$ | 1.13 (1.01—1.26) | $3.40 \times 10^{-2}$ | 1.07 (1.04—1.10) | $2.80 \times 10^{-5}$ | 1.09 (1.04—1.14) | $4.39 \times 10^{-9}$ | 1.09 (1.06—1.12) | $4.39 \times 10^{-9}$ |
| rs75331444 | 2 | 44069772 | G/A | 0.95 | *ABCG8* | Known | 1.22 (0.98—1.52) | $7.50 \times 10^{-2}$ | 0.97 (0.65—1.47) | 0.90 | 1.11 (1.07—1.14) | $2.14 \times 10^{-8}$ | 1.11 (1.08—1.13) | $1.81 \times 10^{-8}$ | 1.11 (1.07—1.14) | $1.81 \times 10^{-8}$ |

| | | | | | | | | | | | | | | | | |
|---|---|---|---|---|---|---|---|---|---|---|---|---|---|---|---|---|
| rs1058588 | 2 | 85808871 | T/C | 0.41 | *VAMP8* | Known | 1.02 (0.98—1.06) | 0.36 | 1.05 (0.97—1.12) | 0.22 | 1.06 (1.04—1.08) | $1.06 \times 10^{-11}$ | 1.05 (1.03—1.07) | $3.52 \times 10^{-11}$ | 1.05 (1.04—1.07) | $3.52 \times 10^{-11}$ |
| rs17678683 | 2 | 145286559 | G/T | 0.08 | *ZEB2 -TEX41* | Known | 1.15 (1.01—1.30) | $3.60 \times 10^{-2}$ | 1.07 (0.88—1.29) | 0.50 | 1.08 (1.05—1.11) | $1.15 \times 10^{-7}$ | 1.08 (1.07—1.10) | $3.72 \times 10^{-8}$ | 1.08 (1.05—1.11) | $3.72 \times 10^{-8}$ |
| rs12619842 | 2 | 164945044 | G/C | 0.79 | *FIGN -GRB14* | Known | 1.08 (1.04—1.12) | $1.79 \times 10^{-4}$ | 1.05 (0.98—1.13) | 0.16 | 1.05 (1.03—1.07) | $8.57 \times 10^{-6}$ | 1.05 (1.04—1.07) | $2.97 \times 10^{-8}$ | 1.05 (1.03—1.08) | $2.97 \times 10^{-8}$ |
| rs72926772 | 2 | 203804289 | T/C | 0.10 | *CARF* | Known | 1.00 (0.80—1.26) | 0.98 | 1.02 (0.98—1.06) | 0.94 | 1.13 (1.10—1.16) | $6.14 \times 10^{-19}$ | 1.13 (1.11—1.15) | $4.68 \times 10^{-18}$ | 1.13 (1.10—1.16) | $4.68 \times 10^{-18}$ |
| rs1250229 | 2 | 216304384 | T/C | 0.26 | *FN1 -MREG* | Known | 1.07 (0.99—1.16) | $7.40 \times 10^{-2}$ | 1.21 (1.05—1.39) | $7.58 \times 10^{-3}$ | 1.07 (1.05—1.09) | $1.85 \times 10^{-13}$ | 1.07 (1.05—1.10) | $1.91 \times 10^{-14}$ | 1.07 (1.05—1.09) | $1.91 \times 10^{-14}$ |
| rs4458205 | 2 | 233557646 | T/C | 0.35 | *EFHD1 -GIGYF2* | Known | 1.09 (1.04—1.15) | $2.80 \times 10^{-4}$ | 1.06 (0.98—1.15) | 0.12 | 1.04 (1.03—1.06) | $3.76 \times 10^{-6}$ | 1.05 (1.03—1.07) | $4.11 \times 10^{-8}$ | 1.05 (1.03—1.07) | $4.11 \times 10^{-8}$ |
| rs293914 | 3 | 14947133 | G/C | 0.13 | *FGD5* | Known | 1.08 (1.04—1.12) | $9.60 \times 10^{-5}$ | 1.08 (1.01—1.16) | $3.20 \times 10^{-2}$ | 1.06 (1.03—1.09) | $9.02 \times 10^{-5}$ | 1.07 (1.06—1.08) | $4.21 \times 10^{-8}$ | 1.07 (1.04—1.09) | $4.21 \times 10^{-8}$ |
| rs11713141 | 3 | 138067626 | C/T | 0.38 | *MRAS* | Known | 1.12 (1.07—1.17) | $1.29 \times 10^{-6}$ | 1.04 (0.97—1.12) | 0.26 | 1.04 (1.03—1.06) | $7.65 \times 10^{-7}$ | 1.05 (1.02—1.08) | $9.94 \times 10^{-10}$ | 1.05 (1.03—1.07) | $9.94 \times 10^{-10}$ |
| rs12645070 | 4 | 57770106 | A/G | 0.23 | *SPINK2 -REST* | Known | 1.07 (1.03—1.12) | $3.94 \times 10^{-4}$ | 0.98 (0.91—1.05) | 0.58 | 1.05 (1.03—1.07) | $3.45 \times 10^{-7}$ | 1.05 (1.02—1.08) | $3.10 \times 10^{-8}$ | 1.05 (1.03—1.07) | $3.10 \times 10^{-8}$ |
| rs11099097 | 4 | 81167309 | T/C | 0.29 | *PRDM8 -FGF5* | Known | 1.05 (1.01—1.09) | $2.40 \times 10^{-2}$ | 1.10 (1.02—1.18) | $1.60 \times 10^{-2}$ | 1.05 (1.03—1.07) | $1.65 \times 10^{-7}$ | 1.05 (1.04—1.06) | $4.15 \times 10^{-9}$ | 1.05 (1.03—1.07) | $4.15 \times 10^{-9}$ |
| rs11099493 | 4 | 82587050 | A/G | 0.70 | *RASGEF1B -HNRNPD* | Known | 1.08 (1.03—1.12) | $9.85 \times 10^{-4}$ | 1.14 (1.05—1.24) | $2.09 \times 10^{-3}$ | 1.04 (1.02—1.06) | $1.47 \times 10^{-5}$ | 1.05 (1.02—1.08) | $4.16 \times 10^{-8}$ | 1.05 (1.03—1.07) | $4.16 \times 10^{-8}$ |
| rs1847331 | 4 | 120936581 | G/T | 0.27 | *PDE5A -MAD2L1* | Known | 1.10 (1.05—1.14) | $1.70 \times 10^{-5}$ | 1.01 (0.93—1.09) | 0.80 | 1.04 (1.03—1.06) | $3.26 \times 10^{-6}$ | 1.05 (1.02—1.08) | $2.15 \times 10^{-8}$ | 1.05 (1.03—1.07) | $2.15 \times 10^{-8}$ |
| rs73855810 | 4 | 148383424 | A/G | 0.17 | *TTC29* | Known | 1.09 | $3.01 \times 10^{-5}$ | 1.08 | $4.10 \times 10^{-2}$ | 1.07 | $4.54 \times 10^{-9}$ | 1.07 | $1.94 \times 10^{-12}$ | 1.07 | $1.94 \times 10^{-12}$ |

| | | | | | | | | | | | | | | | | |
|---|---|---|---|---|---|---|---|---|---|---|---|---|---|---|---|---|
| | | | | | -EDNRA | | (1.05—1.14) | | (1.00—1.17) | | (1.05—1.09) | | (1.06—1.09) | | (1.05—1.09) | |
| rs34914832 | 4 | 156626078 | G/A | 0.79 | GUCY1A3 | Known | 1.08 (1.03—1.13) | $7.99 \times 10^{-4}$ | 1.10 (1.01—1.20) | $2.90 \times 10^{-2}$ | 1.06 (1.04—1.08) | $7.68 \times 10^{-9}$ | 1.07 (1.05—1.08) | $2.56 \times 10^{-11}$ | 1.07 (1.05—1.08) | $2.56 \times 10^{-11}$ |
| rs9381634 | 6 | 13022728 | A/G | 0.30 | PHACTR1 | Known | 1.07 (1.03—1.11) | $5.49 \times 10^{-4}$ | 1.04 (0.97—1.12) | 0.23 | 1.04 (1.03—1.06) | $3.88 \times 10^{-6}$ | 1.05 (1.04—1.06) | $3.31 \times 10^{-8}$ | 1.05 (1.03—1.06) | $3.31 \times 10^{-8}$ |
| rs6456496 | 6 | 22599386 | C/G | 0.33 | HDGFL1 -NRSN1 | Known | 1.05 (1.01—1.10) | $2.40 \times 10^{-2}$ | 1.01 (0.93—1.10) | 0.74 | 1.05 (1.03—1.07) | $1.53 \times 10^{-8}$ | 1.05 (1.04—1.06) | $4.66 \times 10^{-9}$ | 1.05 (1.03—1.07) | $4.66 \times 10^{-9}$ |
| rs3130342 | 6 | 32080146 | C/A | 0.83 | TNXB -ATF6B | Known | 1.10 (1.02—1.18) | $9.10 \times 10^{-3}$ | 1.11 (1.00—1.22) | $4.10 \times 10^{-2}$ | 1.07 (1.05—1.10) | $1.52 \times 10^{-7}$ | 1.08 (1.06—1.09) | $3.09 \times 10^{-9}$ | 1.08 (1.05—1.11) | $3.10 \times 10^{-9}$ |
| rs55902013 | 6 | 39150657 | C/G | 0.80 | SAYSD1 -KCNK5 | Known | 1.12 (1.06—1.18) | $5.92 \times 10^{-5}$ | 1.13 (1.02—1.24) | $2.20 \times 10^{-2}$ | 1.05 (1.03—1.07) | $2.28 \times 10^{-7}$ | 1.06 (1.03—1.09) | $3.85 \times 10^{-10}$ | 1.06 (1.04—1.08) | $3.85 \times 10^{-10}$ |
| rs12193307 | 6 | 134103250 | A/G | 0.66 | TARID | Known | 1.08 (1.04—1.12) | $1.15 \times 10^{-4}$ | 1.08 (1.00—1.16) | $4.00 \times 10^{-2}$ | 1.05 (1.03—1.07) | $9.89 \times 10^{-8}$ | 1.05 (1.04—1.07) | $1.53 \times 10^{-10}$ | 1.05 (1.04—1.07) | $1.53 \times 10^{-10}$ |
| rs56393506 | 6 | 161089307 | T/C | 0.16 | LPA | Known | 1.12 (1.05—1.18) | $4.34 \times 10^{-4}$ | 1.09 (0.98—1.21) | 0.10 | 1.15 (1.12—1.18) | $6.36 \times 10^{-33}$ | 1.15 (1.13—1.17) | $5.12 \times 10^{-34}$ | 1.15 (1.13—1.18) | $5.12 \times 10^{-34}$ |
| rs2107595 | 7 | 19049388 | A/G | 0.20 | HDAC9 -TWIST1 | Known | 1.07 (1.02—1.11) | $1.91 \times 10^{-3}$ | 1.05 (0.97—1.13) | 0.24 | 1.08 (1.06—1.10) | $3.41 \times 10^{-13}$ | 1.07 (1.06—1.08) | $1.34 \times 10^{-14}$ | 1.07 (1.05—1.09) | $1.34 \times 10^{-14}$ |
| rs2107713 | 7 | 107232369 | T/C | 0.30 | BCAP29 | Known | 1.07 (1.03—1.11) | $1.82 \times 10^{-3}$ | 1.06 (0.98—1.15) | 0.12 | 1.04 (1.03—1.06) | $3.14 \times 10^{-6}$ | 1.05 (1.03—1.06) | $4.82 \times 10^{-8}$ | 1.05 (1.03—1.07) | $4.82 \times 10^{-8}$ |
| rs11556924 | 7 | 129663496 | C/T | 0.70 | ZC3HC1 | Known | 1.23 (1.09—1.40) | $1.04 \times 10^{-3}$ | 1.38 (1.09—1.74) | $7.60 \times 10^{-3}$ | 1.07 (1.05—1.09) | $6.26 \times 10^{-13}$ | 1.07 (1.03—1.11) | $4.68 \times 10^{-14}$ | 1.07 (1.05—1.09) | $4.68 \times 10^{-14}$ |
| rs2269997 | 7 | 139723400 | G/A | 0.74 | PARP12 | Known | 1.05 (1.02—1.09) | $6.10 \times 10^{-3}$ | 1.06 (0.98—1.13) | 0.14 | 1.05 (1.03—1.07) | $5.16 \times 10^{-7}$ | 1.05 (1.05—1.05) | $1.31 \times 10^{-8}$ | 1.05 (1.03—1.07) | $1.31 \times 10^{-8}$ |
| rs328 | 8 | 19819724 | C/G | 0.90 | LPL | Known | 1.13 (1.07—1.19) | $5.83 \times 10^{-5}$ | 0.96 (0.86—1.07) | 0.47 | 1.07 (1.04—1.10) | $1.09 \times 10^{-6}$ | 1.07 (1.03—1.12) | $3.35 \times 10^{-8}$ | 1.07 (1.05—1.10) | $3.35 \times 10^{-8}$ |

| | | | | | | | | | | | | | | | | |
|---|---|---|---|---|---|---|---|---|---|---|---|---|---|---|---|---|
| rs2954026 | 8 | 126484526 | T/G | 0.32 | *TRIB1-FAM84B* | Known | 1.04 (1.00—1.09) | $3.20 \times 10^{-2}$ | 1.01 (0.93—1.08) | 0.89 | 1.06 (1.04—1.08) | $6.68 \times 10^{-10}$ | 1.05 (1.04—1.07) | $5.36 \times 10^{-10}$ | 1.05 (1.04—1.07) | $5.36 \times 10^{-10}$ |
| rs10811650 | 9 | 22067593 | G/A | 0.44 | *CDKN2B-AS1* | Known | 1.12 (1.08—1.16) | $4.45 \times 10^{-8}$ | 1.13 (1.05—1.21) | $1.17 \times 10^{-3}$ | 1.15 (1.13—1.17) | $3.21 \times 10^{-67}$ | 1.15 (1.13—1.17) | $1.71 \times 10^{-71}$ | 1.15 (1.13—1.17) | $1.71 \times 10^{-71}$ |
| rs10985343 | 9 | 124415691 | C/T | 0.31 | *DAB2IP* | Known | 1.09 (1.05—1.13) | $3.29 \times 10^{-5}$ | 1.07 (1.00—1.15) | $4.80 \times 10^{-2}$ | 1.04 (1.02—1.06) | $3.53 \times 10^{-5}$ | 1.05 (1.02—1.07) | $4.01 \times 10^{-8}$ | 1.05 (1.03—1.07) | $4.01 \times 10^{-8}$ |
| rs643434 | 9 | 136142355 | A/G | 0.38 | *ABO* | Known | 1.12 (1.07—1.16) | $6.80 \times 10^{-8}$ | 1.12 (1.04—1.20) | $3.14 \times 10^{-3}$ | 1.03 (1.02—1.05) | $3.64 \times 10^{-5}$ | 1.05 (1.01—1.08) | $3.77 \times 10^{-9}$ | 1.05 (1.03—1.06) | $3.77 \times 10^{-9}$ |
| rs10826753 | 10 | 30326776 | C/T | 0.57 | *KIAA1462* | Known | 1.02 (0.98—1.06) | 0.26 | 0.98 (0.91—1.05) | 0.49 | 1.05 (1.04—1.07) | $1.90 \times 10^{-9}$ | 1.04 (1.02—1.07) | $2.19 \times 10^{-8}$ | 1.04 (1.03—1.06) | $2.19 \times 10^{-8}$ |
| rs518594 | 10 | 44757107 | T/C | 0.81 | *ZNF32-CXCL12* | Known | 1.05 (1.01—1.10) | $1.20 \times 10^{-2}$ | 1.11 (1.03—1.19) | $8.26 \times 10^{-3}$ | 1.07 (1.05—1.09) | $7.85 \times 10^{-11}$ | 1.07 (1.06—1.09) | $1.07 \times 10^{-12}$ | 1.07 (1.05—1.09) | $1.07 \times 10^{-12}$ |
| rs2250644 | 10 | 91008879 | T/C | 0.35 | *LIPA* | Known | 1.08 (1.03—1.13) | $1.07 \times 10^{-3}$ | 1.15 (1.06—1.25) | $6.65 \times 10^{-4}$ | 1.06 (1.05—1.08) | $2.97 \times 10^{-13}$ | 1.07 (1.05—1.09) | $3.40 \times 10^{-16}$ | 1.07 (1.05—1.09) | $3.40 \times 10^{-16}$ |
| rs7910601 | 10 | 104583395 | G/T | 0.80 | *WBP1L-CYP17A1* | Known | 1.09 (1.05—1.14) | $5.97 \times 10^{-6}$ | 1.00 (0.93—1.07) | 0.97 | 1.06 (1.04—1.08) | $1.90 \times 10^{-7}$ | 1.06 (1.03—1.09) | $8.74 \times 10^{-10}$ | 1.06 (1.04—1.08) | $8.74 \times 10^{-10}$ |
| rs2257129 | 10 | 122898697 | C/T | 0.90 | *WDR11-FGFR2* | Novel | 1.08 (1.04—1.13) | $3.17 \times 10^{-4}$ | 1.02 (0.95—1.10) | 0.58 | 1.10 (1.06—1.14) | $1.64 \times 10^{-6}$ | 1.08 (1.05—1.12) | $3.95 \times 10^{-8}$ | 1.08 (1.05—1.11) | $3.95 \times 10^{-8}$ |
| rs360153 | 11 | 9762274 | C/T | 0.55 | *SWAP70* | Known | 1.05 (1.01—1.09) | $2.10 \times 10^{-2}$ | 0.95 (0.88—1.02) | 0.16 | 1.05 (1.03—1.06) | $1.39 \times 10^{-8}$ | 1.04 (1.02—1.07) | $3.16 \times 10^{-8}$ | 1.04 (1.03—1.06) | $3.16 \times 10^{-8}$ |
| rs4757137 | 11 | 13291827 | G/T | 0.69 | *RASSF10-ARNTL* | Known | 1.07 (1.03—1.12) | $8.63 \times 10^{-4}$ | 1.03 (0.96—1.11) | 0.35 | 1.04 (1.03—1.06) | $1.64 \times 10^{-6}$ | 1.05 (1.03—1.06) | $3.25 \times 10^{-8}$ | 1.05 (1.03—1.06) | $3.25 \times 10^{-8}$ |
| rs12801636 | 11 | 65391317 | G/A | 0.73 | *PCNXL3* | Known | 1.07 (1.03—1.11) | $2.64 \times 10^{-4}$ | 1.07 (1.00—1.15) | $4.80 \times 10^{-2}$ | 1.04 (1.02—1.06) | $7.75 \times 10^{-6}$ | 1.05 (1.03—1.07) | $1.68 \times 10^{-8}$ | 1.05 (1.03—1.07) | $1.68 \times 10^{-8}$ |
| rs11226018 | 11 | 103664501 | T/A | 0.49 | *DYNC2H1* | Known | 1.14 | $4.81 \times 10\text{-}^{11}$ | 1.07 | $8.90 \times 10^{-2}$ | 1.03 | $2.63 \times 10^{-4}$ | 1.04 | $1.94 \times 10^{-8}$ | 1.04 | $1.94 \times 10^{-8}$ |

| | | | | | | | | | | | | | | | | |
|---|---|---|---|---|---|---|---|---|---|---|---|---|---|---|---|---|
| | | | | | *-PDGFD* | | (1.10—1.18) | | (0.99—1.15) | | (1.01—1.05) | | (1.00—1.09) | | (1.03—1.06) | |
| rs10488763 | 11 | 110244360 | T/A | 0.18 | *RDX-FDX1* | Novel | 1.10 (1.06—1.14) | $2.16 \times 10^{-6}$ | 1.15 (1.07—1.23) | $1.73 \times 10^{-4}$ | 1.05 (1.02—1.07) | $7.96 \times 10^{-5}$ | 1.06 (1.02—1.11) | $9.33 \times 10^{-10}$ | 1.06 (1.04—1.09) | $9.33 \times 10^{-10}$ |
| rs662799 | 11 | 116663707 | G/A | 0.14 | *APOA5* | Known | 1.09 (1.05—1.14) | $1.22 \times 10^{-5}$ | 1.10 (1.02—1.18) | $1.40 \times 10^{-2}$ | 1.06 (1.03—1.09) | $8.19 \times 10^{-5}$ | 1.07 (1.05—1.09) | $6.43 \times 10^{-9}$ | 1.07 (1.04—1.09) | $6.43 \times 10^{-9}$ |
| rs10841443 | 12 | 20220033 | G/C | 0.65 | *LINC02398* | Known | 1.07 (1.03—1.11) | $7.03 \times 10^{-4}$ | 1.05 (0.98—1.13) | 0.19 | 1.04 (1.03—1.06) | $1.03 \times 10^{-6}$ | 1.05 (1.04—1.06) | $1.28 \times 10^{-8}$ | 1.05 (1.03—1.06) | $1.28 \times 10^{-8}$ |
| rs75160195 | 12 | 54521594 | T/C | 0.07 | *HOXC4-SMUG1* | Known | 1.13 (1.07—1.19) | $2.95 \times 10^{-5}$ | 1.05 (0.95—1.15) | 0.36 | 1.09 (1.06—1.13) | $8.64 \times 10^{-7}$ | 1.10 (1.07—1.13) | $2.38 \times 10^{-9}$ | 1.10 (1.06—1.13) | $2.38 \times 10^{-9}$ |
| rs117742247 | 12 | 89991313 | T/C | 0.18 | *ATP2B1* | Known | 1.00 (0.95—1.05) | 0.95 | 1.08 (1.00—1.18) | $6.00 \times 10^{-2}$ | 1.06 (1.04—1.09) | $5.32 \times 10^{-9}$ | 1.06 (1.02—1.09) | $3.07 \times 10^{-8}$ | 1.06 (1.03—1.08) | $3.08 \times 10^{-8}$ |
| rs7311541 | 12 | 95374260 | C/T | 0.90 | *NDUFA12* | Known | 1.14 (1.09—1.19) | $1.36 \times 10^{-8}$ | 1.08 (0.99—1.17) | $9.90 \times 10^{-2}$ | 1.05 (1.02—1.08) | $1.10 \times 10^{-3}$ | 1.07 (1.02—1.13) | $2.47 \times 10^{-8}$ | 1.07 (1.05—1.10) | $2.47 \times 10^{-8}$ |
| rs1558803 | 12 | 109966198 | A/G | 0.02 | *UBE3B* | Known | 1.22 (1.13—1.32) | $5.96 \times 10^{-8}$ | 1.27 (1.12—1.44) | $1.81 \times 10^{-4}$ | 1.13 (1.05—1.22) | $2.12 \times 10^{-3}$ | 1.19 (1.11—1.27) | $1.45 \times 10^{-10}$ | 1.19 (1.12—1.25) | $1.45 \times 10^{-10}$ |
| rs3858704 | 12 | 111705893 | A/G | 0.66 | *CUX2* | Known | 1.26 (1.21—1.31) | $4.66 \times 10^{-30}$ | 1.16 (1.08—1.25) | $5.29 \times 10^{-5}$ | 1.03 (1.01—1.05) | $8.40 \times 10^{-4}$ | 1.06 (0.96—1.17) | $2.57 \times 10^{-13}$ | 1.06 (1.05—1.08) | $2.57 \times 10^{-13}$ |
| rs7139170 | 12 | 113263518 | A/C | 0.67 | *RPH3A* | Known | 1.14 (1.10—1.18) | $9.49 \times 10^{-11}$ | 1.08 (1.00—1.15) | $4.60 \times 10^{-2}$ | 1.04 (1.02—1.06) | $9.86 \times 10^{-4}$ | 1.06 (1.01—1.11) | $6.69 \times 10^{-9}$ | 1.06 (1.04—1.08) | $6.69 \times 10^{-9}$ |
| rs2244608 | 12 | 121416988 | G/A | 0.35 | *HNF1A* | Known | 1.00 (0.96—1.04) | 0.98 | 1.05 (0.98—1.13) | 0.15 | 1.05 (1.03—1.07) | $2.32 \times 10^{-9}$ | 1.05 (1.02—1.07) | $1.99 \times 10^{-8}$ | 1.05 (1.03—1.06) | $1.99 \times 10^{-8}$ |
| rs9513116 | 13 | 29016715 | A/G | 0.33 | *FLT1* | Known | 1.10 (1.05—1.15) | $4.80 \times 10^{-5}$ | 1.04 (0.97—1.12) | 0.25 | 1.04 (1.03—1.06) | $4.39 \times 10^{-7}$ | 1.05 (1.03—1.07) | $2.95 \times 10^{-9}$ | 1.05 (1.03—1.07) | $2.95 \times 10^{-9}$ |
| rs55940034 | 13 | 111043309 | G/A | 0.26 | *COL4A2* | Known | 1.10 (1.03—1.17) | $6.17 \times 10^{-3}$ | 1.07 (0.95—1.22) | 0.26 | 1.06 (1.04—1.07) | $5.16 \times 10^{-9}$ | 1.06 (1.04—1.07) | $4.41 \times 10^{-10}$ | 1.06 (1.04—1.08) | $4.41 \times 10^{-10}$ |

| | | | | | | | | | | | | | | | | |
|---|---|---|---|---|---|---|---|---|---|---|---|---|---|---|---|---|
| rs2098297 | 14 | 75619118 | G/T | 0.51 | *TMED10* | Known | 1.12 (1.06—1.17) | $2.92 \times 10^{-5}$ | 1.07 (0.98—1.18) | 0.14 | 1.04 (1.02—1.06) | $4.15 \times 10^{-6}$ | 1.04 (1.02—1.07) | $4.01 \times 10^{-8}$ | 1.04 (1.03—1.06) | $4.01 \times 10^{-8}$ |
| rs12436072 | 14 | 100135306 | A/G | 0.39 | *HHIPL1* | Known | 1.07 (1.02—1.11) | $3.86 \times 10^{-3}$ | 0.99 (0.92—1.08) | 0.86 | 1.04 (1.03—1.06) | $2.96 \times 10^{-7}$ | 1.04 (1.03—1.06) | $4.71 \times 10^{-8}$ | 1.04 (1.03—1.06) | $4.71 \times 10^{-8}$ |
| rs56062135 | 15 | 67455630 | C/T | 0.81 | *SMAD3* | Known | 1.21 (1.09—1.34) | $1.79 \times 10^{-4}$ | 1.05 (0.88—1.25) | 0.57 | 1.07 (1.05—1.09) | $6.36 \times 10^{-12}$ | 1.08 (1.05—1.11) | $4.73 \times 10^{-13}$ | 1.08 (1.06—1.10) | $4.73 \times 10^{-13}$ |
| rs62010554 | 15 | 79002832 | G/A | 0.25 | *CHRNB4 -ADAMTS7* | Known | 1.04 (0.99—1.08) | $9.20 \times 10^{-2}$ | 1.12 (1.04—1.20) | $2.74 \times 10^{-3}$ | 1.06 (1.04—1.09) | $6.85 \times 10^{-11}$ | 1.06 (1.04—1.08) | $2.83 \times 10^{-12}$ | 1.06 (1.04—1.08) | $2.83 \times 10^{-12}$ |
| rs734780 | 15 | 89564958 | T/C | 0.83 | *MFGE8 -ABHD2* | Known | 1.09 (1.05—1.13) | $4.70 \times 10^{-6}$ | 1.02 (0.95—1.09) | 0.68 | 1.07 (1.04—1.10) | $4.25 \times 10^{-7}$ | 1.07 (1.04—1.10) | $4.38 \times 10^{-10}$ | 1.07 (1.05—1.09) | $4.38 \times 10^{-10}$ |
| rs4932371 | 15 | 91404788 | C/T | 0.32 | *BLM -FURIN* | Known | 1.12 (1.06—1.17) | $3.14 \times 10^{-5}$ | 1.12 (1.01—1.23) | $2.70 \times 10^{-2}$ | 1.05 (1.03—1.07) | $9.49 \times 10^{-9}$ | 1.06 (1.03—1.09) | $1.51 \times 10^{-11}$ | 1.06 (1.04—1.08) | $1.51 \times 10^{-11}$ |
| rs2000999 | 16 | 72108093 | A/G | 0.23 | *HPR* | Known | 1.08 (1.04—1.12) | $2.39 \times 10^{-4}$ | 1.09 (1.02—1.17) | $1.30 \times 10^{-2}$ | 1.06 (1.03—1.08) | $6.71 \times 10^{-7}$ | 1.06 (1.05—1.08) | $4.70 \times 10^{-10}$ | 1.06 (1.04—1.08) | $4.70 \times 10^{-10}$ |
| rs8057203 | 16 | 75404974 | C/T | 0.58 | *CFDP1* | Known | 1.05 (1.01—1.09) | $1.40 \times 10^{-2}$ | 1.10 (1.02—1.18) | $8.71 \times 10^{-3}$ | 1.04 (1.02—1.06) | $2.22 \times 10^{-6}$ | 1.04 (1.03—1.06) | $3.90 \times 10^{-8}$ | 1.04 (1.03—1.06) | $3.90 \times 10^{-8}$ |
| rs2641438 | 17 | 2017786 | G/C | 0.30 | *SMG6* | Known | 1.06 (1.02—1.11) | $7.03 \times 10^{-3}$ | 1.11 (1.02—1.20) | $1.40 \times 10^{-2}$ | 1.05 (1.03—1.07) | $1.67 \times 10^{-7}$ | 1.05 (1.03—1.07) | $2.52 \times 10^{-9}$ | 1.05 (1.03—1.07) | $2.52 \times 10^{-9}$ |
| rs7214227 | 17 | 58991025 | T/C | 0.16 | *BCAS3* | Known | 1.14 (1.08—1.20) | $2.21 \times 10^{-6}$ | 1.11 (1.00—1.22) | $4.40 \times 10^{-2}$ | 1.06 (1.04—1.09) | $1.56 \times 10^{-7}$ | 1.07 (1.04—1.11) | $4.80 \times 10^{-11}$ | 1.07 (1.05—1.10) | $4.80 \times 10^{-11}$ |
| rs2812 | 17 | 62401118 | C/T | 0.54 | *PECAM1* | Known | 1.08 (1.03—1.13) | $3.94 \times 10^{-4}$ | 1.09 (1.01—1.18) | $2.70 \times 10^{-2}$ | 1.04 (1.03—1.06) | $1.77 \times 10^{-6}$ | 1.05 (1.03—1.07) | $1.01 \times 10^{-8}$ | 1.05 (1.03—1.06) | $1.01 \times 10^{-8}$ |
| rs3786723 | 19 | 11166476 | T/C | 0.73 | *SMARCA4* | Known | 1.07 (1.01—1.14) | $1.90 \times 10^{-2}$ | 1.06 (0.95—1.18) | 0.27 | 1.05 (1.03—1.07) | $1.67 \times 10^{-8}$ | 1.05 (1.05—1.06) | $2.19 \times 10^{-9}$ | 1.05 (1.04—1.07) | $2.19 \times 10^{-9}$ |
| rs2241714 | 19 | 41869392 | T/C | 0.35 | *TMEM91* | Known | 1.10 | $4.10 \times 10^{-7}$ | 1.14 | $2.02 \times 10^{-4}$ | 1.04 | $2.55 \times 10^{-6}$ | 1.05 | $3.68 \times 10^{-11}$ | 1.05 | $3.69 \times 10^{-11}$ |

| | | | | | | | | | | | | | | | | |
|---|---|---|---|---|---|---|---|---|---|---|---|---|---|---|---|---|
| | | | | | | | (1.06—1.14) | | (1.06—1.22) | | (1.02—1.06) | | (1.02—1.09) | | (1.04—1.07) | |
| rs56131196 | 19 | 45422846 | A/G | 0.16 | *APOC1* | Known | 1.12 (1.04—1.20) | $1.89 \times 10^{-3}$ | 1.15 (1.02—1.29) | $2.10 \times 10^{-2}$ | 1.09 (1.06—1.11) | $2.71 \times 10^{-12}$ | 1.09 (1.07—1.11) | $2.24 \times 10^{-14}$ | 1.09 (1.07—1.11) | $2.24 \times 10^{-14}$ |
| rs7279974 | 21 | 35666961 | C/G | 0.26 | *SLC5A3* -*KCNE2* | Known | 0.98 (0.89—1.07) | 0.61 | 1.09 (0.93—1.29) | 0.28 | 1.07 (1.05—1.09) | $2.38 \times 10^{-11}$ | 1.06 (1.04—1.09) | $1.24 \times 10^{-10}$ | 1.06 (1.04—1.08) | $1.24 \times 10^{-10}$ |
| rs6004124 | 22 | 24697757 | G/A | 0.95 | *SPECC1L* | Known | 1.13 (1.06—1.20) | $1.66 \times 10^{-4}$ | 1.06 (0.95—1.19) | 0.30 | 1.10 (1.06—1.14) | $7.32 \times 10^{-7}$ | 1.10 (1.08—1.12) | $3.00 \times 10^{-9}$ | 1.10 (1.07—1.14) | $3.00 \times 10^{-9}$ |

Chr., chromosome; A1, effect allele; A2, non-effect allele; freq., frequency; OR, odds ratio; CI, confidence interval; UKBB, UK Biobank.

**Supplemental Table 4. Pleiotropy analysis for newly identified loci**

| Lead variant in new loci | Lead variant in the GWAS catalog data | Selected reasons | $r^2$ in EAS | $r^2$ in EUR | Trait |
|---|---|---|---|---|---|
| rs6587520 | rs2230061 | position | 0.45 | 0.56 | Fat body mass |
| rs6587520 | rs11800059 | LD | 0.99 | 1.00 | Neutrophil percentage of white cells |
| rs6587520 | rs112158604 | position | - | - | Lymphocyte counts |
| rs6587520 | rs11204677 | LD | 0.55 | 0.75 | Monocyte percentage of white cells |
| rs6587520 | rs11204677 | LD | 0.55 | 0.75 | Granulocyte percentage of myeloid white cells |
| rs2257129 | rs1907240 | LD, position | 0.95 | 1.00 | Body mass index |

Pleiotropy analysis using the GWAS catalog data (https://www.ebi.ac.uk/gwas/).

EAS, East Asians; EUR, Europeans; LD, Linkage disequilibrium.

**Supplemental Table 5. Lead variants in novel loci and associated clinical traits**

| Category | Trait | rs6587520 | | | rs2257129 | | | rs10488763 | | |
|---|---|---|---|---|---|---|---|---|---|---|
| | | BETA | SE | FDR | BETA | SE | FDR | BETA | SE | FDR |
| Anthropometric | Body mass index (BMI) | -0.004 | 0.004 | 0.58 | -0.026 | 0.004 | $1.4 \times 10^{-9}$ | -0.001 | 0.004 | 0.91 |
| Metabolic | Total cholesterol (TC) | 0.001 | 0.004 | 0.86 | -0.002 | 0.004 | 0.75 | 0.002 | 0.004 | 0.91 |
| | High-density-lipoprotein cholesterol (HDL-C) | 0.002 | 0.005 | 0.86 | -0.037 | 0.006 | $5.5 \times 10^{-9}$ | -0.002 | 0.005 | 0.91 |
| | Low-density-lipoprotein cholesterol (LDL-C) | 0.003 | 0.005 | 0.73 | 0.001 | 0.006 | 0.89 | 0.010 | 0.005 | 0.42 |
| | Triglyceride (TG) | -0.002 | 0.004 | 0.77 | 0.020 | 0.005 | $4.3 \times 10^{-4}$ | -0.003 | 0.004 | 0.86 |
| | Blood sugar (BS) | 0.008 | 0.005 | 0.34 | 0.007 | 0.005 | 0.39 | 0.005 | 0.005 | 0.81 |
| | Hemoglobin A1c (HbA1c) | 0.004 | 0.007 | 0.77 | 0.018 | 0.007 | 0.12 | -0.001 | 0.007 | 0.96 |
| Protein | Total protein (TP) | 0.017 | 0.004 | $3.0 \times 10^{-3}$ | 0.011 | 0.005 | 0.12 | 0.012 | 0.004 | 0.18 |
| | Albumin (Alb) | 0.001 | 0.004 | 0.86 | 0.010 | 0.005 | 0.21 | -0.002 | 0.005 | 0.91 |
| | Non-albumin protein (NAP) | 0.019 | 0.005 | $2.0 \times 10^{-3}$ | 0.003 | 0.005 | 0.75 | 0.009 | 0.005 | 0.37 |
| | Albumin/globulin ratio (A/G) | -0.015 | 0.005 | $1.6 \times 10^{-2}$ | 0.003 | 0.005 | 0.75 | -0.008 | 0.005 | 0.42 |
| Kidney-related | Blood urea nitrogen (BUN) | 0.000 | 0.004 | 0.97 | 0.011 | 0.004 | $5.8 \times 10^{-2}$ | -0.004 | 0.004 | 0.81 |
| | Serum creatinine (sCr) | 0.004 | 0.004 | 0.58 | 0.002 | 0.004 | 0.78 | -0.004 | 0.004 | 0.81 |
| | Estimated glomerular filtration rate (eGFR) | -0.005 | 0.004 | 0.58 | -0.002 | 0.004 | 0.75 | 0.004 | 0.004 | 0.81 |

| | | | | | | | | | | |
|---|---|---|---|---|---|---|---|---|---|---|
| | Uric acid (UA) | 0.005 | 0.004 | 0.58 | 0.000 | 0.005 | 0.97 | 0.000 | 0.004 | 0.96 |
| Electrolyte | Sodium (Na) | -0.007 | 0.004 | 0.40 | 0.006 | 0.004 | 0.39 | -0.005 | 0.004 | 0.81 |
| | Potassium (K) | -0.001 | 0.004 | 0.92 | 0.003 | 0.004 | 0.75 | -0.006 | 0.004 | 0.61 |
| | Chloride (Cl) | -0.004 | 0.004 | 0.58 | 0.006 | 0.004 | 0.39 | -0.002 | 0.004 | 0.91 |
| | Calcium (Ca) | 0.006 | 0.005 | 0.58 | 0.008 | 0.006 | 0.39 | 0.013 | 0.005 | 0.29 |
| | Phosphorus (P) | -0.007 | 0.007 | 0.58 | 0.003 | 0.007 | 0.78 | -0.007 | 0.007 | 0.81 |
| Liver-related | Total bilirubin (TBil) | 0.001 | 0.004 | 0.86 | 0.000 | 0.005 | 0.97 | -0.004 | 0.004 | 0.85 |
| | Zinc sulfate turbidity test (ZTT) | 0.009 | 0.013 | 0.73 | -0.015 | 0.014 | 0.48 | 0.009 | 0.013 | 0.85 |
| | Aspartate aminotransferase (AST) | 0.014 | 0.004 | $5.0 \times 10^{-3}$ | -0.001 | 0.004 | 0.83 | 0.005 | 0.004 | 0.71 |
| | Alanine aminotransferase (ALT) | 0.006 | 0.004 | 0.41 | 0.005 | 0.004 | 0.48 | 0.001 | 0.004 | 0.91 |
| | Alkaline phosphatase (ALP) | 0.005 | 0.004 | 0.58 | -0.002 | 0.005 | 0.78 | 0.005 | 0.004 | 0.81 |
| | γ-glutamyl transferase (GGT) | 0.005 | 0.004 | 0.58 | 0.007 | 0.004 | 0.36 | -0.001 | 0.004 | 0.91 |
| Other biochemical | Activated partial thromboplastin time (APTT) | 0.002 | 0.007 | 0.86 | 0.021 | 0.008 | $8.9 \times 10^{-2}$ | 0.001 | 0.007 | 0.96 |
| | Prothrombin time (PT) | 0.012 | 0.006 | 0.25 | -0.008 | 0.006 | 0.44 | 0.004 | 0.006 | 0.85 |
| | Fibrinogen (Fbg) | 0.011 | 0.011 | 0.60 | 0.021 | 0.011 | 0.26 | 0.026 | 0.011 | 0.29 |
| | Creatine kinase (CK) | 0.013 | 0.004 | $2.6 \times 10^{-2}$ | 0.010 | 0.005 | 0.20 | 0.000 | 0.004 | 0.98 |
| | Lactate dehydrogenase (LDH) | 0.003 | 0.004 | 0.73 | 0.004 | 0.004 | 0.52 | 0.002 | 0.004 | 0.91 |

| | | | | | | | | | | |
|---|---|---|---|---|---|---|---|---|---|---|
| | C-reactive protein (CRP) | 0.003 | 0.005 | 0.73 | 0.005 | 0.006 | 0.57 | 0.016 | 0.005 | 0.18 |
| | White blood cell count (WBC) | 0.005 | 0.004 | 0.58 | 0.002 | 0.005 | 0.81 | 0.001 | 0.004 | 0.91 |
| | Neutrophil count (Neutro) | 0.017 | 0.006 | $2.7 \times 10^{-2}$ | 0.001 | 0.006 | 0.93 | 0.008 | 0.006 | 0.73 |
| | Eosinophil count (Eosino) | 0.005 | 0.006 | 0.69 | 0.009 | 0.006 | 0.39 | 0.001 | 0.006 | 0.92 |
| | Basophil count (Baso) | 0.016 | 0.006 | $3.1 \times 10^{-2}$ | -0.003 | 0.006 | 0.75 | -0.003 | 0.006 | 0.91 |
| | Monocyte count (Mono) | -0.015 | 0.006 | $6.8 \times 10^{-2}$ | 0.007 | 0.006 | 0.50 | -0.003 | 0.006 | 0.91 |
| | Lymphocyte count (Lym) | -0.004 | 0.006 | 0.73 | 0.013 | 0.006 | 0.20 | 0.004 | 0.006 | 0.85 |
| | Red blood cell count (RBC) | 0.002 | 0.004 | 0.77 | 0.005 | 0.005 | 0.48 | 0.003 | 0.004 | 0.85 |
| | Hemoglobin (Hb) | 0.004 | 0.004 | 0.67 | -0.001 | 0.005 | 0.88 | 0.002 | 0.004 | 0.91 |
| | Hematocrit (Ht) | 0.002 | 0.004 | 0.77 | -0.003 | 0.005 | 0.75 | 0.001 | 0.004 | 0.91 |
| | Mean corpuscular volume (MCV) | 0.004 | 0.004 | 0.62 | -0.014 | 0.005 | $4.6 \times 10^{-2}$ | -0.003 | 0.004 | 0.85 |
| | Mean corpuscular hemoglobin (MCH) | 0.005 | 0.004 | 0.58 | -0.006 | 0.005 | 0.46 | -0.001 | 0.004 | 0.92 |
| | Mean corpuscular hemoglobin concentration (MCHC) | 0.004 | 0.004 | 0.61 | 0.007 | 0.005 | 0.39 | 0.004 | 0.004 | 0.81 |
| | Platelet count (Plt) | 0.000 | 0.004 | 0.93 | -0.004 | 0.005 | 0.57 | 0.003 | 0.004 | 0.85 |
| Blood pressure | Systolic blood pressure (SBP) | 0.009 | 0.004 | 0.12 | 0.008 | 0.004 | 0.22 | 0.008 | 0.004 | 0.39 |
| | Diastolic blood pressure (DBP) | -0.001 | 0.004 | 0.92 | 0.007 | 0.004 | 0.36 | 0.003 | 0.004 | 0.85 |
| | Mean arterial pressure (MAP) | 0.004 | 0.004 | 0.61 | 0.008 | 0.004 | 0.22 | 0.005 | 0.004 | 0.73 |

| | | | | | | | | | | |
|---|---|---|---|---|---|---|---|---|---|---|
| | Pulse pressure (PP) | 0.014 | 0.004 | $6.0 \times 10^{-3}$ | 0.006 | 0.004 | 0.39 | 0.008 | 0.004 | 0.37 |
| Echocardiographic | Interventricular septum thickness (IVS) | 0.007 | 0.010 | 0.73 | 0.018 | 0.011 | 0.36 | -0.003 | 0.010 | 0.91 |
| | Posterior wall thickness (PW) | -0.005 | 0.010 | 0.80 | 0.013 | 0.011 | 0.46 | 0.008 | 0.010 | 0.85 |
| | Left ventricular internal dimension in diastole (LVDd) | 0.020 | 0.010 | 0.25 | 0.002 | 0.011 | 0.90 | 0.007 | 0.010 | 0.85 |
| | Left ventricular internal dimension in systole (LVDs) | 0.011 | 0.010 | 0.58 | 0.007 | 0.011 | 0.75 | 0.015 | 0.010 | 0.71 |
| | Left ventricular mass (LVM) | 0.016 | 0.010 | 0.42 | 0.016 | 0.011 | 0.39 | 0.006 | 0.010 | 0.90 |
| | Left ventricular mass index (LVMI) | 0.015 | 0.011 | 0.56 | 0.021 | 0.012 | 0.29 | 0.004 | 0.011 | 0.91 |
| | Relative wall thickness (RWT) | -0.016 | 0.010 | 0.44 | 0.005 | 0.011 | 0.78 | 0.001 | 0.010 | 0.96 |
| | Fractional shortening (FS) | 0.003 | 0.010 | 0.86 | -0.008 | 0.011 | 0.73 | -0.021 | 0.010 | 0.37 |
| | Ejection fraction (EF) | 0.005 | 0.010 | 0.77 | -0.014 | 0.011 | 0.42 | -0.012 | 0.010 | 0.81 |
| | E/A ratio (E/A) | 0.002 | 0.015 | 0.92 | -0.007 | 0.017 | 0.78 | 0.003 | 0.015 | 0.93 |

Association between clinical traits and lead variants in novel loci. Clinical traits with FDR < 0.05 are printed in red. FDR was calculated using the Benjamini-Hochberg procedure.

BETA, beta coefficient; SE, standard error; FDR, false discovery rate.

**Supplemental Table 6. Biological pathways suggested by MAGENTA**

| Database | Gene Set | Exp. Genes above 95% cutoff | Obs. Genes above 95% cutoff | GSEA *P*-value 95% cutoff | FDR 95% cutoff | Category |
|---|---|---|---|---|---|---|
| GOTERM | reverse cholesterol transport | 1 | 7 | $9.90 \times 10^{-7}$ | $1.00 \times 10^{-4}$ | lipid/transport |
| GOTERM | cholesterol transporter activity | 1 | 6 | $8.00 \times 10^{-6}$ | $6.50 \times 10^{-4}$ | lipid/transport |
| Ingenuity | LXR.RXR.Activation | 2 | 9 | $2.50 \times 10^{-5}$ | $1.60 \times 10^{-3}$ | lipid/inflammation |
| GOTERM | high-density lipoprotein particle remodeling | 1 | 5 | $1.00 \times 10^{-4}$ | $5.47 \times 10^{-3}$ | lipid |
| GOTERM | cholesterol homeostasis | 2 | 10 | $4.00 \times 10^{-6}$ | $5.83 \times 10^{-3}$ | lipid/homeostasis |
| GOTERM | low-density lipoprotein receptor binding | 1 | 5 | $1.00 \times 10^{-4}$ | $6.04 \times 10^{-3}$ | lipid/binding |
| Ingenuity | Actin.Cytoskeleton.Signaling | 1 | 7 | $2.00 \times 10^{-4}$ | $7.60 \times 10^{-3}$ | signaling |
| PANTHER BIOLOGICAL PROCESS | Tumor_suppressor | 4 | 13 | $2.70 \times 10^{-5}$ | $1.16 \times 10^{-2}$ | cell cycle |
| GOTERM | triglyceride metabolic process | 1 | 7 | $1.00 \times 10^{-4}$ | $1.16 \times 10^{-2}$ | lipid |
| GOTERM | chylomicron | 0 | 4 | $6.00 \times 10^{-4}$ | $1.28 \times 10^{-2}$ | lipid |
| GOTERM | synaptic transmission, cholinergic | 1 | 5 | $4.00 \times 10^{-4}$ | $1.39 \times 10^{-2}$ | signaling |

| | | | | | | |
|---|---|---|---|---|---|---|
| GOTERM | glutathione biosynthetic process | 0 | 4 | $4.00 \times 10^{-4}$ | $1.66 \times 10^{-2}$ | metabolic |
| GOTERM | vasodilation | 1 | 5 | $5.00 \times 10^{-4}$ | $2.06 \times 10^{-2}$ | blood vessel |
| GOTERM | low-density lipoprotein particle | 1 | 4 | $7.00 \times 10^{-4}$ | $2.09 \times 10^{-2}$ | lipid |
| GOTERM | cellular response to starvation | 1 | 4 | $1.00 \times 10^{-3}$ | $2.57 \times 10^{-2}$ | cellular response |
| GOTERM | cholesterol efflux | 1 | 5 | $1.00 \times 10^{-3}$ | $3.19 \times 10^{-2}$ | lipid/transport |
| GOTERM | lipid homeostasis | 1 | 4 | $1.10 \times 10^{-3}$ | $3.91 \times 10^{-2}$ | lipid/homeostasis |
| GOTERM | cholesterol metabolic process | 3 | 10 | $1.00 \times 10^{-4}$ | $4.01 \times 10^{-2}$ | lipid |
| Ingenuity | Neurotrophin.TRK.Signaling | 1 | 4 | $4.90 \times 10^{-3}$ | $4.12 \times 10^{-2}$ | signaling |
| GOTERM | positive regulation of endothelial cell proliferation | 1 | 6 | $7.00 \times 10^{-4}$ | $4.46 \times 10^{-2}$ | cell proliferation |
| GOTERM | negative regulation of mitotic cell cycle | 1 | 4 | $1.80 \times 10^{-3}$ | $4.52 \times 10^{-2}$ | cell cycle |
| GOTERM | microtubule binding | 3 | 10 | $2.75 \times 10^{-4}$ | $4.63 \times 10^{-2}$ | binding |
| GOTERM | cellular response to hormone stimulus | 1 | 5 | $1.80 \times 10^{-3}$ | $4.71 \times 10^{-2}$ | cellular response |
| GOTERM | phospholipid efflux | 0 | 3 | $4.50 \times 10^{-3}$ | $4.71 \times 10^{-2}$ | lipid/transport |
| GOTERM | collagen binding | 2 | 7 | $3.00 \times 10^{-4}$ | $4.77 \times 10^{-2}$ | binding |

Exp, expected; Obs, observed; GSEA, Gene Set Enrichment Analysis.

**Supplemental Table 7. Comparison of risk allele frequencies between EAS and EUR in 76 lead variants**

| SNP | Chr. | Position (GRCh37) | A1/A2 | A1 Freq in EAS* | A1 Freq in EUR* | SNP | Chr. | Position (GRCh37) | A1/A2 | A1 Freq in EAS* | A1 Freq in EUR* |
|---|---|---|---|---|---|---|---|---|---|---|---|
| **Japanese-frequent loci** | | | | | | **European-frequent loci** | | | | | |
| rs11206510 | 1 | 55496039 | T/C | 0.955 | 0.828 | rs6587520 | 1 | 150709723 | T/C | 0.461 | 0.476 |
| rs56348932 | 1 | 56988551 | A/C | 0.979 | 0.882 | rs11810571 | 1 | 151762308 | G/C | 0.506 | 0.860 |
| rs4970837 | 1 | 109822008 | T/G | 0.950 | 0.665 | rs3008629 | 1 | 222835930 | T/C | 0.723 | 0.843 |
| rs12129500 | 1 | 154423764 | T/C | 0.572 | 0.451 | rs11124924 | 2 | 21206275 | G/C | 0.847 | 0.918 |
| rs16986953 | 2 | 19942473 | A/G | 0.347 | 0.089 | rs1058588 | 2 | 85808871 | T/C | 0.386 | 0.431 |
| rs75331444 | 2 | 44069772 | G/A | 0.985 | 0.921 | rs17678683 | 2 | 145286559 | G/T | 0.057 | 0.087 |
| rs4458205 | 2 | 233557646 | T/C | 0.343 | 0.285 | rs12619842 | 2 | 164945044 | G/C | 0.596 | 0.837 |
| rs293914 | 3 | 14947133 | G/C | 0.382 | 0.053 | rs72926772 | 2 | 203804289 | T/C | 0.016 | 0.130 |
| rs12645070 | 4 | 57770106 | A/G | 0.379 | 0.186 | rs1250229 | 2 | 216304384 | T/C | 0.077 | 0.232 |
| rs11099097 | 4 | 81167309 | T/C | 0.359 | 0.265 | rs11713141 | 3 | 138067626 | C/T | 0.272 | 0.333 |
| rs11099493 | 4 | 82587050 | A/G | 0.785 | 0.646 | rs1847331 | 4 | 120936581 | G/T | 0.154 | 0.275 |
| rs73855810 | 4 | 148383424 | A/G | 0.229 | 0.145 | rs6456496 | 6 | 22599386 | C/G | 0.189 | 0.363 |
| rs34914832 | 4 | 156626078 | G/A | 0.768 | 0.766 | rs12193307 | 6 | 134103250 | A/G | 0.601 | 0.674 |

| | | | | | | | | | | | |
|---|---|---|---|---|---|---|---|---|---|---|---|
| rs9381634 | 6 | 13022728 | A/G | 0.505 | 0.260 | rs56393506 | 6 | 161089307 | T/C | 0.094 | 0.182 |
| rs3130342 | 6 | 32080146 | C/A | 0.918 | 0.893 | rs2107713 | 7 | 107232369 | T/C | 0.317 | 0.323 |
| rs55902013 | 6 | 39150657 | C/G | 0.844 | 0.770 | rs2269997 | 7 | 139723400 | G/A | 0.472 | 0.788 |
| rs2107595 | 7 | 19049388 | A/G | 0.358 | 0.168 | rs2954026 | 8 | 126484526 | T/G | 0.272 | 0.318 |
| rs11556924 | 7 | 129663496 | C/T | 0.950 | 0.623 | rs643434 | 9 | 136142355 | A/G | 0.381 | 0.400 |
| rs328 | 8 | 19819724 | C/G | 0.878 | 0.870 | rs518594 | 10 | 44757107 | T/C | 0.658 | 0.854 |
| rs10811650 | 9 | 22067593 | G/A | 0.526 | 0.440 | rs2250644 | 10 | 91008879 | T/C | 0.330 | 0.358 |
| rs10985343 | 9 | 124415691 | C/T | 0.457 | 0.232 | rs7910601 | 10 | 104583395 | G/T | 0.581 | 0.882 |
| rs10826753 | 10 | 30326776 | C/T | 0.659 | 0.552 | rs2257129 | 10 | 122898697 | C/T | 0.613 | 0.968 |
| rs11226018 | 11 | 103664501 | T/A | 0.632 | 0.449 | rs360153 | 11 | 9762274 | C/T | 0.383 | 0.597 |
| rs10488763 | 11 | 110244360 | T/A | 0.372 | 0.113 | rs4757137 | 11 | 13291827 | G/T | 0.564 | 0.754 |
| rs662799 | 11 | 116663707 | G/A | 0.288 | 0.084 | rs12801636 | 11 | 65391317 | G/A | 0.573 | 0.776 |
| rs75160195 | 12 | 54521594 | T/C | 0.199 | 0.046 | rs10841443 | 12 | 20220033 | G/C | 0.594 | 0.673 |
| rs117742247 | 12 | 89991313 | T/C | 0.210 | 0.137 | rs7311541 | 12 | 95374260 | C/T | 0.850 | 0.941 |
| rs1558803 | 12 | 109966198 | A/G | 0.083 | 0.009 | rs3858704 | 12 | 111705893 | A/G | 0.275 | 0.729 |
| rs2244608 | 12 | 121416988 | G/A | 0.392 | 0.342 | rs7139170 | 12 | 113263518 | A/C | 0.502 | 0.715 |
| rs9513116 | 13 | 29016715 | A/G | 0.398 | 0.343 | rs55940034 | 13 | 111043309 | G/A | 0.099 | 0.309 |

| | | | | | | | | | | | |
|---|---|---|---|---|---|---|---|---|---|---|---|
| rs2098297 | 14 | 75619118 | G/T | 0.839 | 0.504 | rs12436072 | 14 | 100135306 | A/G | 0.260 | 0.425 |
| rs56062135 | 15 | 67455630 | C/T | 0.972 | 0.785 | rs734780 | 15 | 89564958 | T/C | 0.531 | 0.889 |
| rs62010554 | 15 | 79002832 | G/A | 0.401 | 0.213 | rs4932371 | 15 | 91404788 | C/T | 0.170 | 0.359 |
| rs2000999 | 16 | 72108093 | A/G | 0.310 | 0.192 | rs8057203 | 16 | 75404974 | C/T | 0.548 | 0.573 |
| rs2812 | 17 | 62401118 | C/T | 0.687 | 0.563 | rs2641438 | 17 | 2017786 | G/C | 0.260 | 0.292 |
| rs3786723 | 19 | 11166476 | T/C | 0.902 | 0.704 | rs7214227 | 17 | 58991025 | T/C | 0.121 | 0.160 |
| rs2241714 | 19 | 41869392 | T/C | 0.554 | 0.325 | rs56131196 | 19 | 45422846 | A/G | 0.099 | 0.198 |
| | | | | | | rs7279974 | 21 | 35666961 | C/G | 0.057 | 0.258 |
| | | | | | | rs6004124 | 22 | 24697757 | G/A | 0.915 | 0.973 |

Chr., chromosome; A1, risk allele; A2, non-risk allele; freq., frequency; EAS, East Asians; EUR, Europeans.

* 1000 Genomes phase 3 (http://www.internationalgenome.org/category/phase-3/)

**Supplemental Table 8. Comparison of risk allele frequencies between EAS and EUR in 230 suggestive significant variants**

| SNP | Chr. | Position (GRCh37) | A1/A2 | A1 Freq in EAS* | A1 Freq in EUR* | SNP | Chr. | Position (GRCh37) | A1/A2 | A1 Freq in EAS* | A1 Freq in EUR* |
|---|---|---|---|---|---|---|---|---|---|---|---|
| **Japanese-frequent loci** | | | | | | **European-frequent loci** | | | | | |
| rs150183244 | 1 | 51356091 | G/T | 0.922 | 0.903 | rs12743493 | 1 | 2224836 | G/A | 0.568 | 0.626 |
| rs34232196 | 1 | 55489542 | C/T | 0.855 | 0.766 | rs2651925 | 1 | 3101863 | C/G | 0.708 | 0.764 |
| rs17111652 | 1 | 55590465 | T/C | 0.051 | 0.040 | rs28380108 | 1 | 38409103 | C/G | 0.317 | 0.425 |
| rs287230 | 1 | 55660292 | G/C | 0.844 | 0.833 | rs11206803 | 1 | 56877509 | T/C | 0.443 | 0.528 |
| rs72664355 | 1 | 57007791 | T/C | 0.978 | 0.884 | rs6587520 | 1 | 150709723 | T/C | 0.461 | 0.476 |
| rs7528419 | 1 | 109817192 | A/G | 0.956 | 0.787 | rs11810571 | 1 | 151762308 | G/C | 0.506 | 0.860 |
| rs11552449 | 1 | 114448389 | T/C | 0.587 | 0.191 | rs6413828 | 1 | 175329712 | A/T | 0.295 | 0.321 |
| rs6689306 | 1 | 154395946 | A/G | 0.563 | 0.442 | rs2820315 | 1 | 201872264 | T/C | 0.100 | 0.287 |
| rs16986953 | 2 | 19942473 | A/G | 0.347 | 0.088 | rs77815753 | 1 | 222730939 | C/T | 0.706 | 0.980 |
| rs7571647 | 2 | 21476634 | G/A | 0.946 | 0.878 | rs67180937 | 1 | 222823743 | G/T | 0.582 | 0.738 |
| rs12469758 | 2 | 21533107 | A/T | 0.054 | 0.023 | rs2854725 | 2 | 21237786 | T/G | 0.851 | 0.912 |
| rs76866386 | 2 | 44075483 | T/C | 0.982 | 0.921 | rs13306194 | 2 | 21252534 | G/A | 0.872 | 0.998 |
| rs2161967 | 2 | 218680529 | T/G | 0.527 | 0.446 | rs58953077 | 2 | 21510295 | T/C | 0.328 | 0.387 |

| | | | | | | | | | | | |
|---|---|---|---|---|---|---|---|---|---|---|---|
| rs2972146 | 2 | 227100698 | T/G | 0.933 | 0.625 | rs6738009 | 2 | 85737509 | A/C | 0.396 | 0.488 |
| rs11691714 | 2 | 230012076 | T/A | 0.572 | 0.572 | rs17678683 | 2 | 145286559 | G/T | 0.057 | 0.086 |
| rs13003675 | 2 | 233584109 | T/C | 0.353 | 0.285 | rs12691704 | 2 | 145822778 | A/G | 0.029 | 0.350 |
| rs35462537 | 3 | 14899394 | A/G | 0.705 | 0.510 | rs12619842 | 2 | 164945044 | G/C | 0.596 | 0.837 |
| rs67913572 | 3 | 14925876 | A/G | 0.180 | 0.028 | rs114123510 | 2 | 203831212 | A/T | 0.016 | 0.130 |
| rs293910 | 3 | 14949102 | T/G | 0.381 | 0.053 | rs10174652 | 2 | 204427340 | G/A | 0.095 | 0.134 |
| rs9811982 | 3 | 49624377 | A/C | 0.929 | 0.667 | rs1250229 | 2 | 216304384 | T/C | 0.077 | 0.232 |
| rs12497018 | 3 | 50301571 | A/G | 0.618 | 0.011 | rs4678145 | 3 | 124450081 | C/G | 0.055 | 0.129 |
| rs13324341 | 3 | 138070901 | T/C | 0.239 | 0.151 | rs2699425 | 4 | 3478051 | C/T | 0.324 | 0.339 |
| rs10513507 | 3 | 156983742 | C/T | 0.549 | 0.351 | rs7678555 | 4 | 120909501 | C/A | 0.159 | 0.311 |
| rs17081933 | 4 | 57822869 | A/T | 0.361 | 0.185 | rs4593108 | 4 | 148281001 | C/G | 0.672 | 0.844 |
| rs10857147 | 4 | 81181072 | T/A | 0.353 | 0.264 | rs2306556 | 4 | 156638573 | A/G | 0.787 | 0.795 |
| rs11099493 | 4 | 82587050 | A/G | 0.785 | 0.646 | rs869396 | 4 | 169688000 | C/A | 0.434 | 0.529 |
| rs28590383 | 4 | 146803248 | T/C | 0.575 | 0.557 | rs4074793 | 5 | 52193125 | G/A | 0.069 | 0.078 |
| rs10305838 | 4 | 148400256 | C/T | 0.227 | 0.146 | rs459193 | 5 | 55806751 | G/A | 0.488 | 0.709 |
| rs2880099 | 4 | 156432368 | A/C | 0.510 | 0.115 | rs28650790 | 5 | 55861464 | T/C | 0.136 | 0.178 |
| rs1842896 | 4 | 156511459 | T/G | 0.730 | 0.530 | rs251023 | 5 | 140893410 | G/A | 0.219 | 0.336 |

| | | | | | | | | | | | |
|---|---|---|---|---|---|---|---|---|---|---|---|
| rs10512704 | 5 | 4048803 | T/C | 0.813 | 0.691 | rs216140 | 5 | 149443298 | T/C | 0.537 | 0.723 |
| rs12916 | 5 | 74656539 | C/T | 0.511 | 0.413 | rs10948377 | 6 | 12987237 | A/G | 0.681 | 0.702 |
| rs1800449 | 5 | 121413208 | T/C | 0.188 | 0.162 | rs6908574 | 6 | 22610165 | C/T | 0.188 | 0.364 |
| rs2081914 | 5 | 122673622 | C/G | 0.752 | 0.475 | rs733701 | 6 | 39171862 | T/C | 0.105 | 0.236 |
| rs7454157 | 6 | 12909874 | G/A | 0.919 | 0.613 | rs1330633 | 6 | 57148971 | G/A | 0.031 | 0.074 |
| rs3130342 | 6 | 32080146 | C/A | 0.918 | 0.893 | rs117665398 | 6 | 82450442 | C/T | 0.126 | 0.165 |
| rs12211281 | 6 | 39147179 | C/T | 0.844 | 0.770 | rs12202017 | 6 | 134173151 | A/G | 0.503 | 0.707 |
| rs4711750 | 6 | 43757082 | A/T | 0.579 | 0.521 | rs577849 | 6 | 149615163 | C/G | 0.634 | 0.873 |
| rs1931656 | 6 | 82610188 | A/T | 0.485 | 0.445 | rs9456496 | 6 | 160443390 | G/A | 0.019 | 0.102 |
| rs11153071 | 6 | 97039741 | G/A | 0.964 | 0.792 | rs41269133 | 6 | 161087863 | T/C | 0.765 | 0.892 |
| rs2327433 | 6 | 134214227 | G/A | 0.181 | 0.098 | rs56393506 | 6 | 161089307 | T/C | 0.094 | 0.182 |
| rs1810126 | 6 | 160872151 | T/C | 0.461 | 0.353 | rs2314852 | 6 | 161122787 | A/G | 0.361 | 0.807 |
| rs9457927 | 6 | 160910282 | G/A | 0.022 | 0.010 | rs2520251 | 7 | 107148928 | A/T | 0.316 | 0.321 |
| rs6956990 | 7 | 5724091 | C/T | 0.024 | 0.012 | rs6945612 | 7 | 139728046 | T/C | 0.461 | 0.787 |
| rs2107595 | 7 | 19049388 | A/G | 0.358 | 0.168 | rs1892609 | 8 | 37380139 | T/C | 0.503 | 0.963 |
| rs59261051 | 7 | 99650429 | T/C | 0.406 | 0.274 | rs302953 | 8 | 106512481 | T/C | 0.426 | 0.572 |
| rs11556924 | 7 | 129663496 | C/T | 0.950 | 0.623 | rs2954029 | 8 | 126490972 | A/T | 0.448 | 0.552 |

| | | | | | | | | | | | |
|---|---|---|---|---|---|---|---|---|---|---|---|
| rs11780610 | 8 | 18259876 | C/T | 0.704 | 0.282 | rs1970014 | 9 | 110517130 | C/A | 0.209 | 0.283 |
| rs2083636 | 8 | 19865263 | T/G | 0.765 | 0.715 | rs781622 | 9 | 114927849 | T/C | 0.339 | 0.365 |
| rs6989064 | 8 | 19941448 | C/T | 0.813 | 0.423 | rs3891088 | 9 | 136083695 | C/T | 0.468 | 0.596 |
| rs2891168 | 9 | 22098619 | G/A | 0.531 | 0.492 | rs1887318 | 10 | 30321598 | T/C | 0.193 | 0.434 |
| rs7045889 | 9 | 22133251 | A/G | 0.722 | 0.606 | rs623932 | 10 | 30479752 | T/A | 0.717 | 0.745 |
| rs4149311 | 9 | 107588777 | T/C | 0.685 | 0.121 | rs640577 | 10 | 44764219 | T/C | 0.659 | 0.854 |
| rs10818583 | 9 | 124422261 | A/G | 0.283 | 0.215 | rs11000448 | 10 | 74682633 | T/G | 0.694 | 0.948 |
| rs9411476 | 9 | 136127601 | A/G | 0.153 | 0.007 | rs2271271 | 10 | 75558867 | A/G | 0.235 | 0.757 |
| rs507666 | 9 | 136149399 | A/G | 0.190 | 0.186 | rs2270552 | 10 | 75863750 | T/C | 0.342 | 0.729 |
| rs11257613 | 10 | 12284392 | G/A | 0.674 | 0.491 | rs7098414 | 10 | 82214586 | A/C | 0.020 | 0.266 |
| rs1870634 | 10 | 44480811 | G/T | 0.756 | 0.666 | rs2246942 | 10 | 91004886 | G/A | 0.332 | 0.361 |
| rs2281674 | 10 | 124130308 | C/G | 0.131 | 0.074 | rs55917128 | 10 | 100023359 | T/C | 0.249 | 0.551 |
| rs2300432 | 10 | 124243189 | C/T | 0.358 | 0.185 | rs11191425 | 10 | 104625970 | C/T | 0.711 | 0.908 |
| rs56210063 | 11 | 8789165 | C/G | 0.186 | 0.060 | rs79780963 | 10 | 104952499 | C/T | 0.021 | 0.034 |
| rs10767932 | 11 | 32382641 | G/A | 0.728 | 0.476 | rs2257129 | 10 | 122898697 | C/T | 0.613 | 0.968 |
| rs2727020 | 11 | 49111407 | C/G | 0.761 | 0.675 | rs360157 | 11 | 9754221 | T/C | 0.370 | 0.590 |
| rs12146487 | 11 | 64026639 | G/A | 0.943 | 0.845 | rs3993105 | 11 | 13303071 | T/C | 0.563 | 0.726 |

| | | | | | | | | | | | |
|---|---|---|---|---|---|---|---|---|---|---|---|
| rs2839812 | 11 | 103673294 | T/A | 0.596 | 0.272 | rs12801636 | 11 | 65391317 | G/A | 0.573 | 0.776 |
| rs10488763 | 11 | 110244360 | T/A | 0.372 | 0.113 | rs11235604 | 11 | 72533536 | C/T | 0.900 | 1.000 |
| rs662799 | 11 | 116663707 | G/A | 0.288 | 0.083 | rs571353 | 11 | 75152243 | C/T | 0.076 | 0.285 |
| rs75160195 | 12 | 54521594 | T/C | 0.199 | 0.046 | rs634552 | 11 | 75282052 | G/T | 0.554 | 0.836 |
| rs2681472 | 12 | 90008959 | G/A | 0.316 | 0.145 | rs3133293 | 11 | 77195100 | G/T | 0.604 | 0.705 |
| rs115704395 | 12 | 109746991 | A/G | 0.154 | 0.000 | rs12221736 | 11 | 110271577 | C/T | 0.770 | 0.868 |
| rs12230500 | 12 | 109850791 | G/A | 0.169 | 0.119 | rs567040 | 11 | 111460678 | C/T | 0.054 | 0.349 |
| rs11067475 | 12 | 110046648 | A/G | 0.081 | 0.009 | rs3962568 | 12 | 20223017 | A/C | 0.495 | 0.700 |
| rs2040571 | 12 | 111357727 | A/G | 0.202 | 0.085 | rs4842680 | 12 | 90116362 | A/G | 0.366 | 0.563 |
| rs4766519 | 12 | 111370980 | C/T | 0.507 | 0.488 | rs12817989 | 12 | 95512389 | T/A | 0.851 | 0.942 |
| rs117741012 | 12 | 111827203 | A/G | 0.495 | 0.003 | rs916682 | 12 | 111699146 | A/G | 0.286 | 0.756 |
| rs12231744 | 12 | 112477055 | C/T | 0.574 | 0.003 | rs7139170 | 12 | 113263518 | A/C | 0.502 | 0.715 |
| rs11066165 | 12 | 112542651 | A/C | 0.574 | 0.040 | rs4766660 | 12 | 113276736 | A/G | 0.406 | 0.588 |
| rs10850001 | 12 | 112553032 | A/T | 0.575 | 0.446 | rs11057830 | 12 | 125307053 | A/G | 0.095 | 0.165 |
| rs1045670 | 12 | 113335681 | A/G | 0.300 | 0.060 | rs117468552 | 13 | 89435862 | C/G | 0.947 | 0.989 |
| rs2244608 | 12 | 121416988 | G/A | 0.392 | 0.342 | rs3783113 | 13 | 110834746 | T/C | 0.353 | 0.354 |
| rs11057401 | 12 | 124427306 | T/A | 0.921 | 0.651 | rs9521678 | 13 | 110916118 | T/C | 0.504 | 0.745 |

| | | | | | | | | | | | |
|---|---|---|---|---|---|---|---|---|---|---|---|
| rs9508027 | 13 | 28986781 | A/C | 0.505 | 0.309 | rs11619113 | 13 | 110918660 | G/C | 0.010 | 0.125 |
| rs9515203 | 13 | 111049623 | T/C | 0.887 | 0.766 | rs4773141 | 13 | 110954353 | G/C | 0.279 | 0.381 |
| rs10143132 | 14 | 75525850 | A/G | 0.837 | 0.503 | rs10139550 | 14 | 100145710 | G/C | 0.360 | 0.435 |
| rs72743461 | 15 | 67441750 | C/A | 0.972 | 0.785 | rs7164479 | 15 | 79123054 | T/C | 0.503 | 0.577 |
| rs62010525 | 15 | 78968310 | G/C | 0.418 | 0.177 | rs4778730 | 15 | 79253321 | T/G | 0.605 | 0.954 |
| rs17581137 | 15 | 96146414 | A/C | 0.917 | 0.778 | rs2351254 | 15 | 89580299 | C/G | 0.559 | 0.893 |
| rs247616 | 16 | 56989590 | C/T | 0.825 | 0.708 | rs2071382 | 15 | 91428197 | T/C | 0.189 | 0.471 |
| rs2000999 | 16 | 72108093 | A/G | 0.310 | 0.192 | rs9930506 | 16 | 53830465 | G/A | 0.218 | 0.438 |
| rs2925979 | 16 | 81534790 | T/C | 0.417 | 0.288 | rs8052763 | 16 | 75251659 | C/G | 0.781 | 0.857 |
| rs12602492 | 17 | 1984121 | C/G | 0.274 | 0.154 | rs3851738 | 16 | 75387533 | C/G | 0.545 | 0.574 |
| rs9897596 | 17 | 17593453 | T/C | 0.828 | 0.505 | rs7500448 | 16 | 83045790 | A/G | 0.732 | 0.756 |
| rs8068844 | 17 | 40571284 | C/T | 0.350 | 0.313 | rs7209460 | 17 | 2048713 | T/C | 0.403 | 0.727 |
| rs4968248 | 17 | 44993128 | G/A | 0.418 | 0.401 | rs1122326 | 17 | 40274873 | C/A | 0.166 | 0.233 |
| rs11655587 | 17 | 47140794 | C/T | 0.837 | 0.681 | rs112502960 | 17 | 47439302 | A/G | 0.216 | 0.364 |
| rs8067638 | 17 | 58930921 | T/C | 0.101 | 0.003 | rs8079832 | 17 | 47513548 | C/T | 0.205 | 0.379 |
| rs6504218 | 17 | 62408299 | G/A | 0.686 | 0.565 | rs1867250 | 17 | 59004233 | A/T | 0.123 | 0.161 |
| rs35489971 | 17 | 72700943 | A/G | 0.265 | 0.181 | rs2068151 | 18 | 32700478 | G/A | 0.777 | 0.889 |

| | | | | | | | | | | | |
|---|---|---|---|---|---|---|---|---|---|---|---|
| rs4393627 | 17 | 73887996 | A/G | 0.908 | 0.901 | rs7229491 | 18 | 46516424 | C/G | 0.444 | 0.670 |
| rs3813126 | 18 | 20037576 | A/G | 0.579 | 0.271 | rs663129 | 18 | 57838401 | A/G | 0.183 | 0.240 |
| rs7251746 | 19 | 10665951 | G/A | 0.890 | 0.839 | rs56307353 | 19 | 1946185 | T/G | 0.292 | 0.563 |
| rs17698352 | 19 | 10909914 | A/G | 0.332 | 0.069 | rs73015715 | 19 | 17855840 | T/C | 0.019 | 0.229 |
| rs12979495 | 19 | 10964632 | G/A | 0.833 | 0.729 | rs8105002 | 19 | 21919934 | T/A | 0.774 | 0.859 |
| rs1433099 | 19 | 11242658 | C/T | 0.744 | 0.708 | rs11881953 | 19 | 41808249 | G/A | 0.725 | 0.829 |
| rs33823 | 19 | 34000725 | C/T | 0.498 | 0.359 | rs7412 | 19 | 45412079 | C/T | 0.900 | 0.937 |
| rs1982072 | 19 | 41864509 | T/A | 0.558 | 0.311 | rs56131196 | 19 | 45422846 | A/G | 0.099 | 0.198 |
| rs7256920 | 19 | 46203083 | G/A | 0.570 | 0.496 | rs34761425 | 19 | 46170711 | T/A | 0.858 | 0.939 |
| rs13734 | 20 | 17594729 | A/G | 0.529 | 0.158 | rs17122844 | 20 | 33452600 | C/T | 0.703 | 0.810 |
| rs6102322 | 20 | 39872768 | T/C | 0.770 | 0.294 | rs259983 | 20 | 57735457 | C/A | 0.063 | 0.160 |
| rs56313611 | 20 | 47456856 | C/T | 0.964 | 0.854 | rs7277800 | 21 | 35655734 | A/G | 0.056 | 0.250 |
| rs555715 | 20 | 48198739 | G/A | 0.377 | 0.231 | rs8140812 | 22 | 24553535 | G/A | 0.926 | 0.939 |
| rs3813452 | 20 | 61174357 | T/C | 0.663 | 0.371 | rs180803 | 22 | 24658858 | G/T | 0.920 | 0.986 |
| rs1999323 | 21 | 30534128 | T/C | 0.189 | 0.140 | rs5760389 | 22 | 24784051 | A/T | 0.921 | 0.984 |
| rs2834414 | 21 | 35598406 | C/T | 0.454 | 0.319 | rs9608859 | 22 | 30667277 | C/T | 0.569 | 0.572 |
| rs5760271 | 22 | 24634471 | T/C | 0.535 | 0.485 | rs4450 | 22 | 33278904 | T/A | 0.152 | 0.583 |

Chr., chromosome; A1, risk allele; A2, non-risk allele; freq., frequency; EAS, East Asians; EUR, Europeans.

* 1000 Genomes phase 3 (http://www.internationalgenome.org/category/phase-3/)

**Supplemental Table 9. Population-specific tissue/cell type enrichment analysis by DEPICT**

| Name | MeSH first level term | META | | East Asians | | Europeans | |
|---|---|---|---|---|---|---|---|
| | | *P* value | FDR < 5% | *P* value | FDR < 5% | *P* value | FDR < 5% |
| Arteries | Cardiovascular System | $3.05 \times 10^{-6}$ | Yes | $3.28 \times 10^{-2}$ | No | $4.99 \times 10^{-4}$ | Yes |
| Adrenal Glands | Endocrine System | $1.31 \times 10^{-5}$ | Yes | $1.02 \times 10^{-3}$ | Yes | $8.09 \times 10^{-3}$ | No |
| Adipose Tissue | Tissues | $1.78 \times 10^{-5}$ | Yes | $6.65 \times 10^{-3}$ | No | $1.43 \times 10^{-3}$ | Yes |
| Adrenal Cortex | Endocrine System | $2.85 \times 10^{-5}$ | Yes | $2.11 \times 10^{-3}$ | Yes | $1.03 \times 10^{-2}$ | No |
| Subcutaneous Fat | Tissues | $3.13 \times 10^{-5}$ | Yes | $6.81 \times 10^{-3}$ | No | $2.16 \times 10^{-3}$ | Yes |
| Adipose Tissue, White | Tissues | $3.13 \times 10^{-5}$ | Yes | $6.81 \times 10^{-3}$ | No | $2.16 \times 10^{-3}$ | Yes |
| Subcutaneous Fat, Abdominal | Tissues | $3.84 \times 10^{-5}$ | Yes | $8.79 \times 10^{-3}$ | No | $2.37 \times 10^{-3}$ | Yes |
| Abdominal Fat | Tissues | $3.84 \times 10^{-5}$ | Yes | $8.79 \times 10^{-3}$ | No | $2.37 \times 10^{-3}$ | Yes |
| Adipocytes | Cells | $1.88 \times 10^{-4}$ | Yes | $2.77 \times 10^{-2}$ | No | $8.88 \times 10^{-3}$ | No |
| Stomach | Digestive System | $2.27 \times 10^{-4}$ | Yes | $2.10 \times 10^{-2}$ | No | $1.74 \times 10^{-3}$ | Yes |
| Upper Gastrointestinal Tract | Digestive System | $2.86 \times 10^{-4}$ | Yes | $4.22 \times 10^{-2}$ | No | $1.74 \times 10^{-3}$ | Yes |
| Connective Tissue Cells | Cells | $4.81 \times 10^{-4}$ | Yes | 0.15 | No | $9.06 \times 10^{-2}$ | No |
| Blood Vessels | Cardiovascular System | $7.94 \times 10^{-4}$ | Yes | 0.16 | No | $1.15 \times 10^{-2}$ | No |
| Cecum | Digestive System | $9.63 \times 10^{-4}$ | Yes | $3.44 \times 10^{-2}$ | No | $3.87 \times 10^{-3}$ | Yes |

| | | | | | | | |
|---|---|---|---|---|---|---|---|
| Serous Membrane | Tissues | $1.83 \times 10^{-3}$ | Yes | $5.19 \times 10^{-3}$ | No | $2.21 \times 10^{-2}$ | No |
| Muscle, Smooth | Tissues | $1.85 \times 10^{-3}$ | Yes | 0.28 | No | $2.07 \times 10^{-2}$ | No |
| Ileum | Digestive System | $2.10 \times 10^{-3}$ | Yes | $5.86 \times 10^{-3}$ | No | $4.04 \times 10^{-2}$ | No |
| Intestine, Small | Digestive System | $2.63 \times 10^{-3}$ | Yes | $7.21 \times 10^{-3}$ | No | $2.32 \times 10^{-2}$ | No |
| Joints | Musculoskeletal System | $2.85 \times 10^{-3}$ | Yes | 0.12 | No | $2.81 \times 10^{-2}$ | No |
| Joint Capsule | Musculoskeletal System | $2.85 \times 10^{-3}$ | Yes | 0.12 | No | $2.81 \times 10^{-2}$ | No |
| Synovial Membrane | Musculoskeletal System | $2.85 \times 10^{-3}$ | Yes | 0.12 | No | $2.81 \times 10^{-2}$ | No |
| Liver | Digestive System | $4.00 \times 10^{-3}$ | Yes | $3.72 \times 10^{-2}$ | No | $1.60 \times 10^{-2}$ | No |
| Muscle Cells | Cells | $4.42 \times 10^{-3}$ | Yes | 0.29 | No | $6.68 \times 10^{-2}$ | No |
| Myocytes, Smooth Muscle | Cells | $4.42 \times 10^{-3}$ | Yes | 0.29 | No | $6.68 \times 10^{-2}$ | No |
| Myometrium | Urogenital System | $4.73 \times 10^{-3}$ | Yes | 0.17 | No | $8.08 \times 10^{-3}$ | No |
| Lower Gastrointestinal Tract | Digestive System | $5.10 \times 10^{-3}$ | Yes | $2.84 \times 10^{-2}$ | No | $1.96 \times 10^{-2}$ | No |
| Hepatocytes | Cells | $5.15 \times 10^{-3}$ | Yes | 0.18 | No | $1.69 \times 10^{-2}$ | No |
| Rectum | Digestive System | $5.19 \times 10^{-3}$ | Yes | $3.17 \times 10^{-2}$ | No | $1.49 \times 10^{-2}$ | No |
| Intestine, Large | Digestive System | $6.81 \times 10^{-3}$ | Yes | $3.37 \times 10^{-2}$ | No | $2.21 \times 10^{-2}$ | No |
| Colon | Digestive System | $8.23 \times 10^{-3}$ | Yes | $3.60 \times 10^{-2}$ | No | $2.50 \times 10^{-2}$ | No |
| Veins | Cardiovascular System | $8.71 \times 10^{-3}$ | Yes | 0.20 | No | $4.02 \times 10^{-2}$ | No |

DEPICT, Data-driven Expression-Prioritized Integration for Complex Traits.

MeSH, Medical Subjects Heading. FDR, false discovery rare.

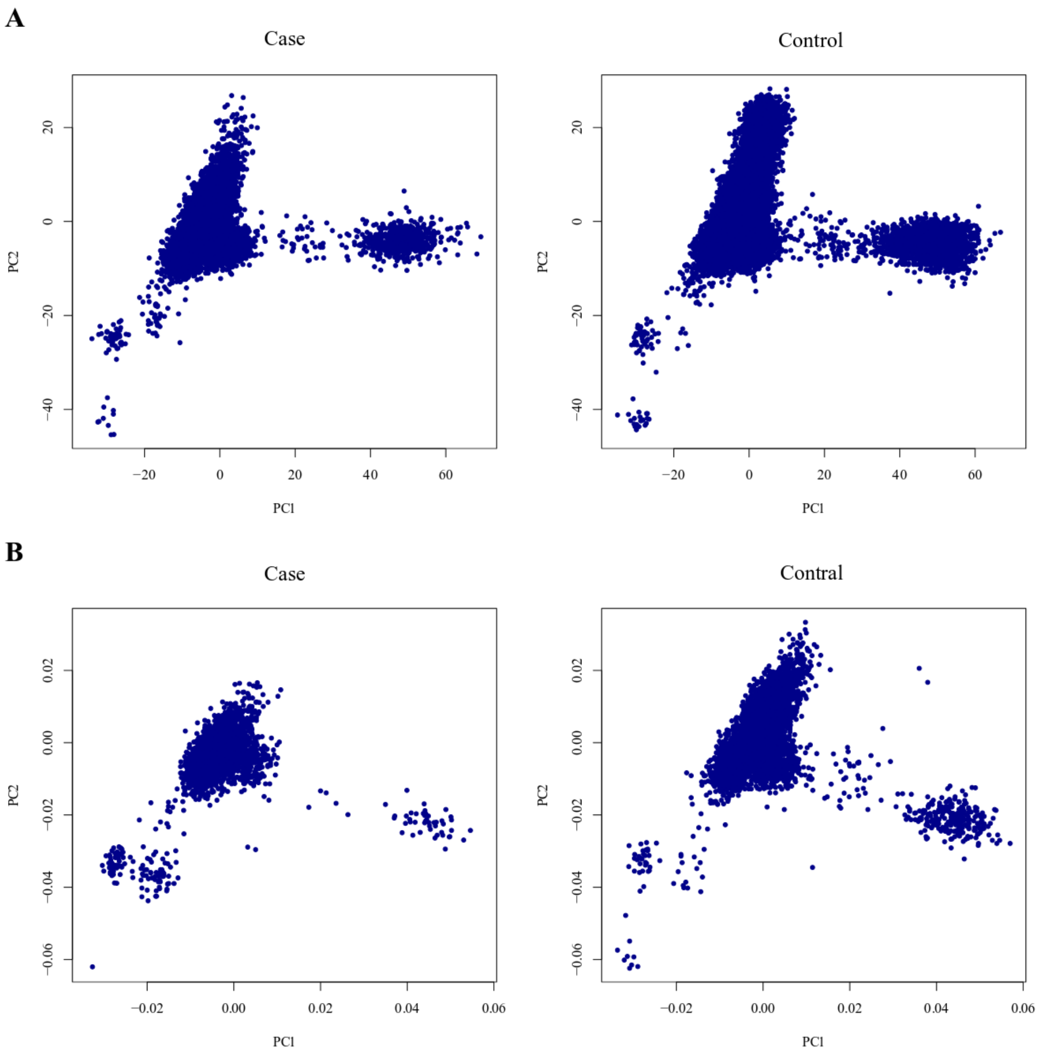


**Supplemental Figure 1. Principal component analysis (PCA) in Japanese GWAS**

**A**, PCA in the first Japanese GWAS. In the first study, all cases were selected from BioBank Japan (BBJ).

**B**, PCA in the second Japanese GWAS. In the second study, all cases were selected from Osaka Acute Coronary Insufficiency Study (OACIS).

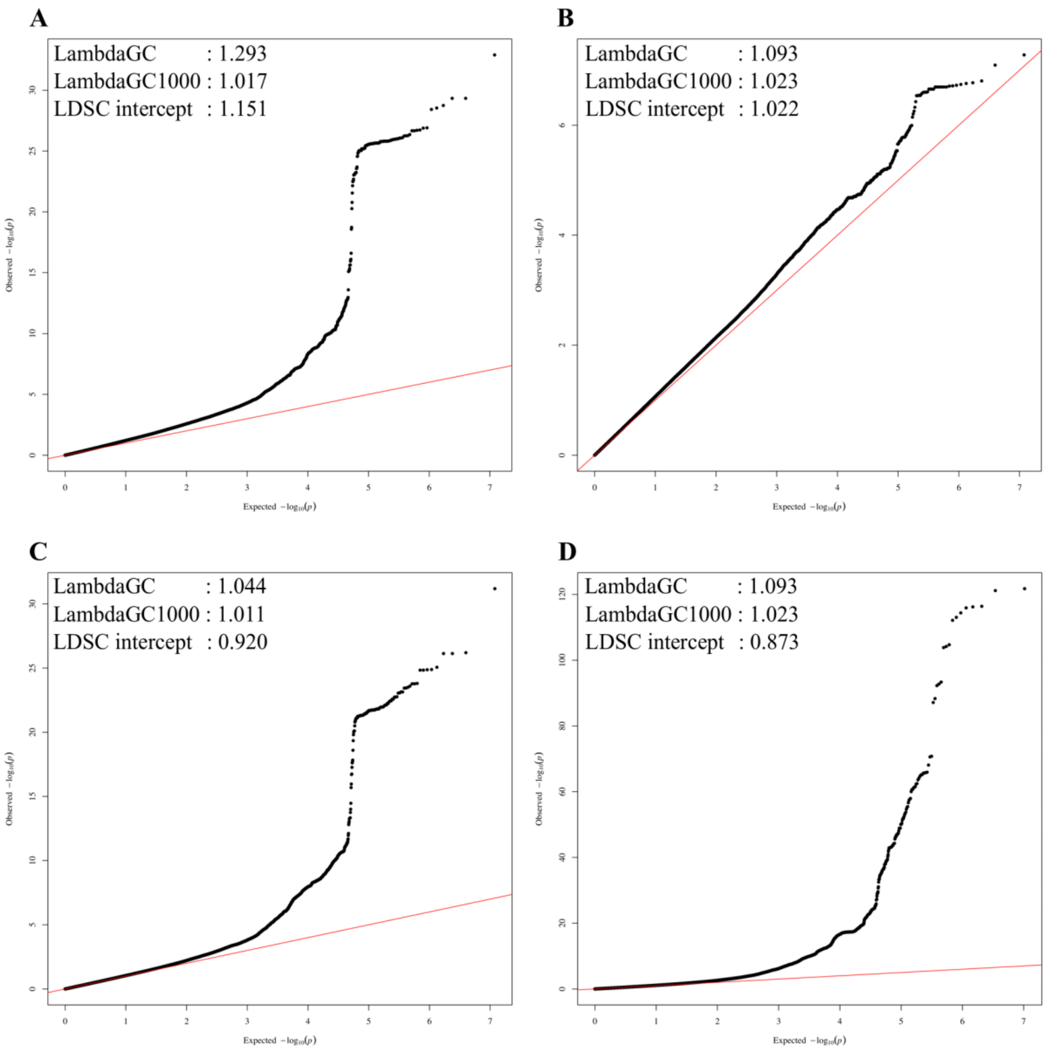


**Supplemental Figure 2. Q-Q plots in Japanese Study 1, 2, meta-analysis and trans-ethnic meta-analysis**

The figure shows quantile-quantile plot for **A**, the first Japanese GWAS, **B**, the second Japanese GWAS, **C**, the Japanese meta-GWAS, and **D**, Transethnic meta-analysis. LDSC, linkage disequilibrium score regression

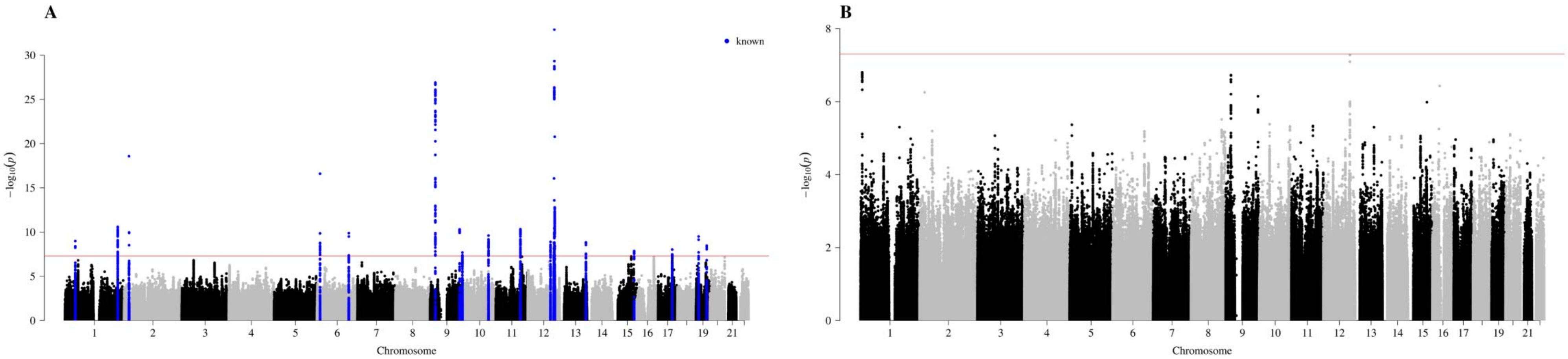


**Supplemental Figure 3. Manhattan plots in the first and second Japanese GWAS**

Manhattan plots for **A**, The first Japanese GWAS (12,494 CAD cases and 28,879 controls) **B**, The second Japanese GWAS (2,808 CAD cases and 7,261 controls). The *x* axis denotes chromosomal location and the *y* axis denotes $-\log_{10} P$ value for each SNP. Blue dots represent known GWS loci (lead variants ± 250 kb). The red horizontal line shows the threshold of GWS ($P = 5.0 \times 10^{-8}$).

**A**

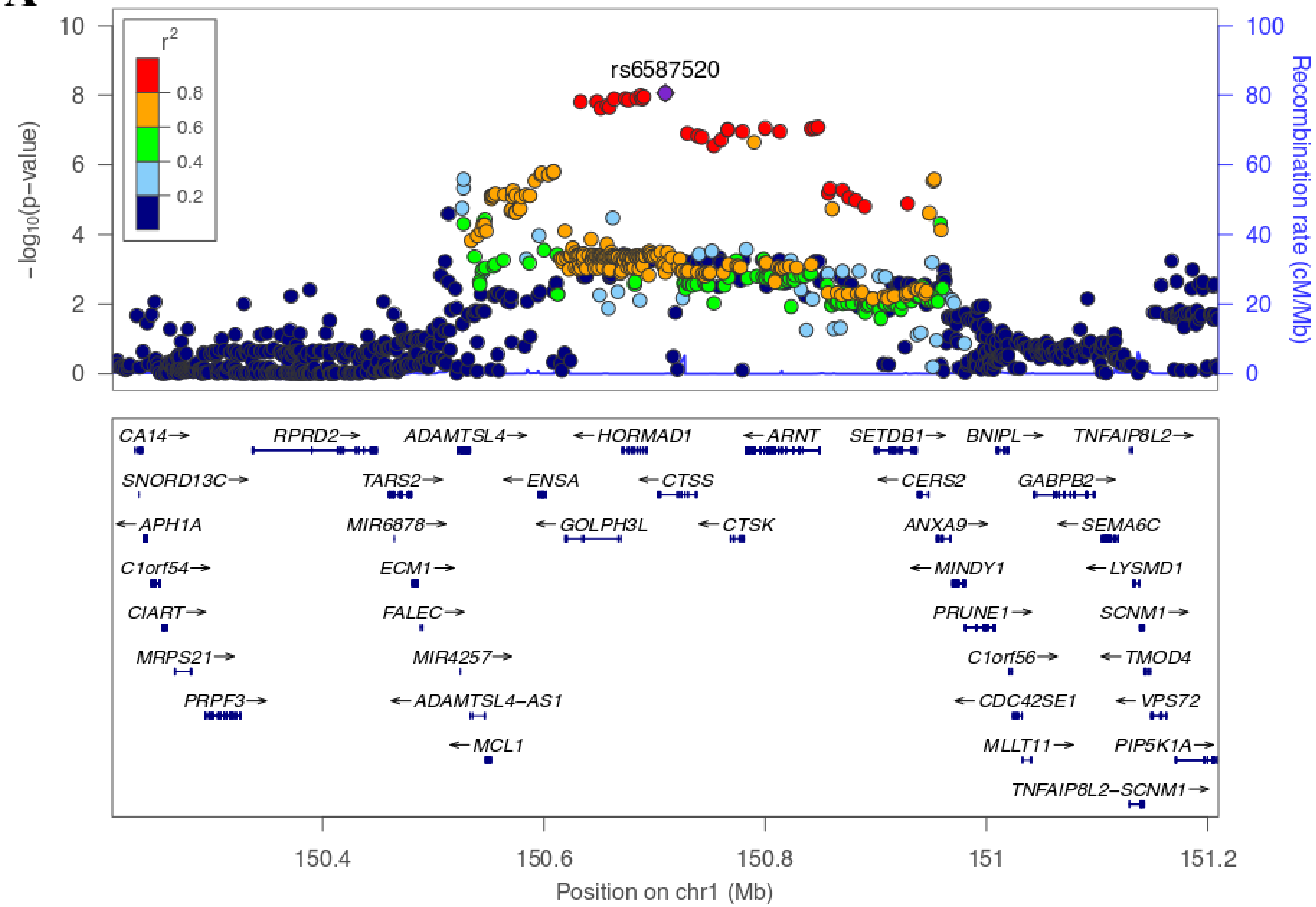

rs6587520
r2
0.8
0.6
0.4
0.2
−log10(p−value)
Recombination rate (cM/Mb)
CA14 RPRD2 ADAMTSL4 HORMAD1 ARNT SETDB1 BNIPL TNFAIP8L2
SNORD13C TARS2 ENSA CTSS CERS2 GABPB2
APH1A MIR6878 GOLPH3L CTSK ANXA9 SEMA6C
C1orf54 ECM1 MINDY1 LYSMD1
CIART FALEC PRUNE1 SCNM1
MRPS21 MIR4257 C1orf56 TMOD4
PRPF3 ADAMTSL4−AS1 CDC42SE1 VPS72
MCL1 MLLT11 PIP5K1A
TNFAIP8L2−SCNM1
150.4 150.6 150.8 151 151.2
Position on chr1 (Mb)

B

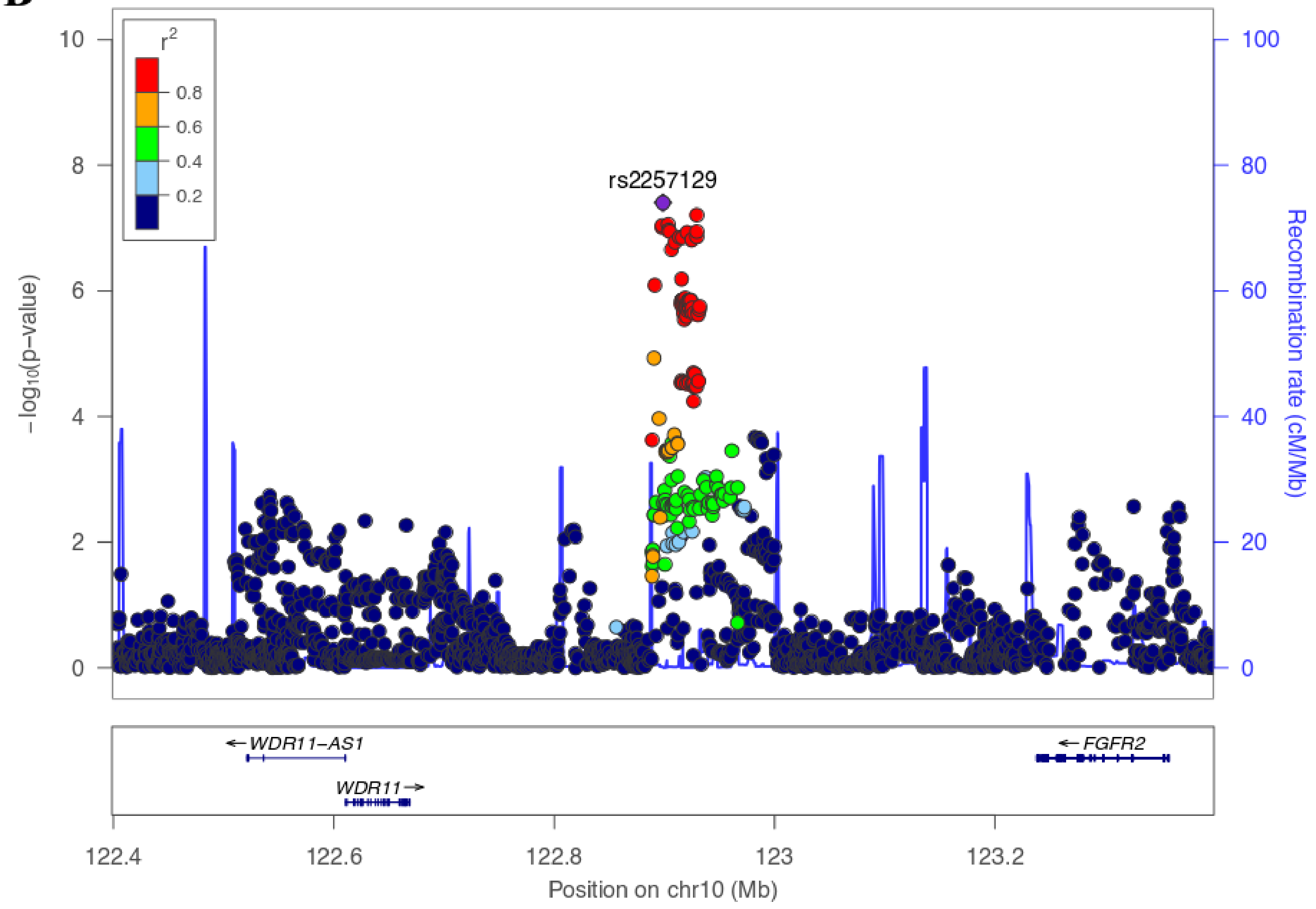

r2
0.8
0.6
0.4
0.2
rs2257129
10
8
6
4
2
0
−log10(p−value)
100
80
60
40
20
0
Recombination rate (cM/Mb)
←WDR11−AS1
WDR11→
←FGFR2
122.4
122.6
122.8
123
123.2
Position on chr10 (Mb)

C

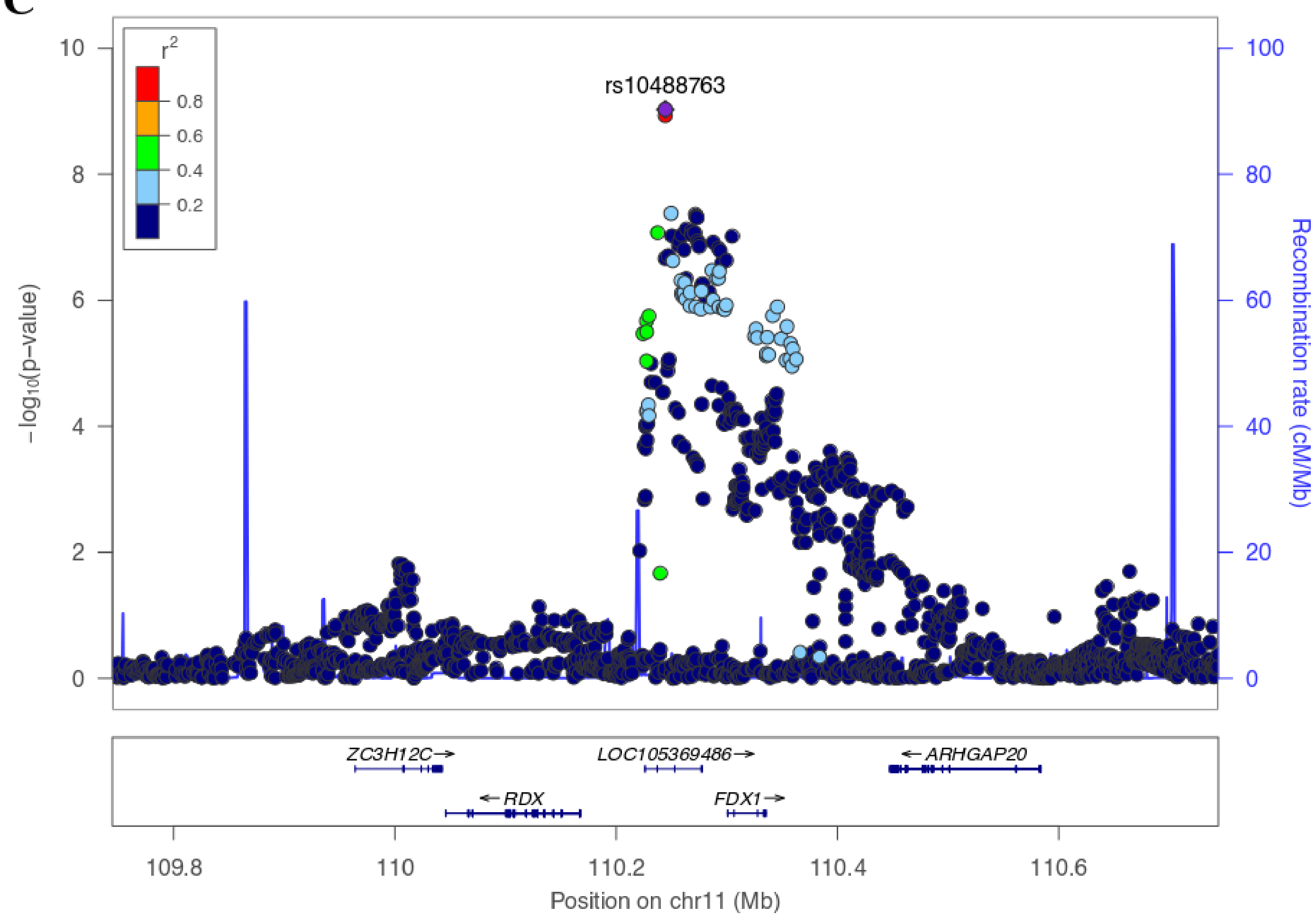

rs10488763
r2
0.8
0.6
0.4
0.2
−log10(p−value)
Recombination rate (cM/Mb)
10
8
6
4
2
0
100
80
60
40
20
0
ZC3H12C
LOC105369486
ARHGAP20
RDX
FDX1
109.8
110
110.2
110.4
110.6
Position on chr11 (Mb)

**D**

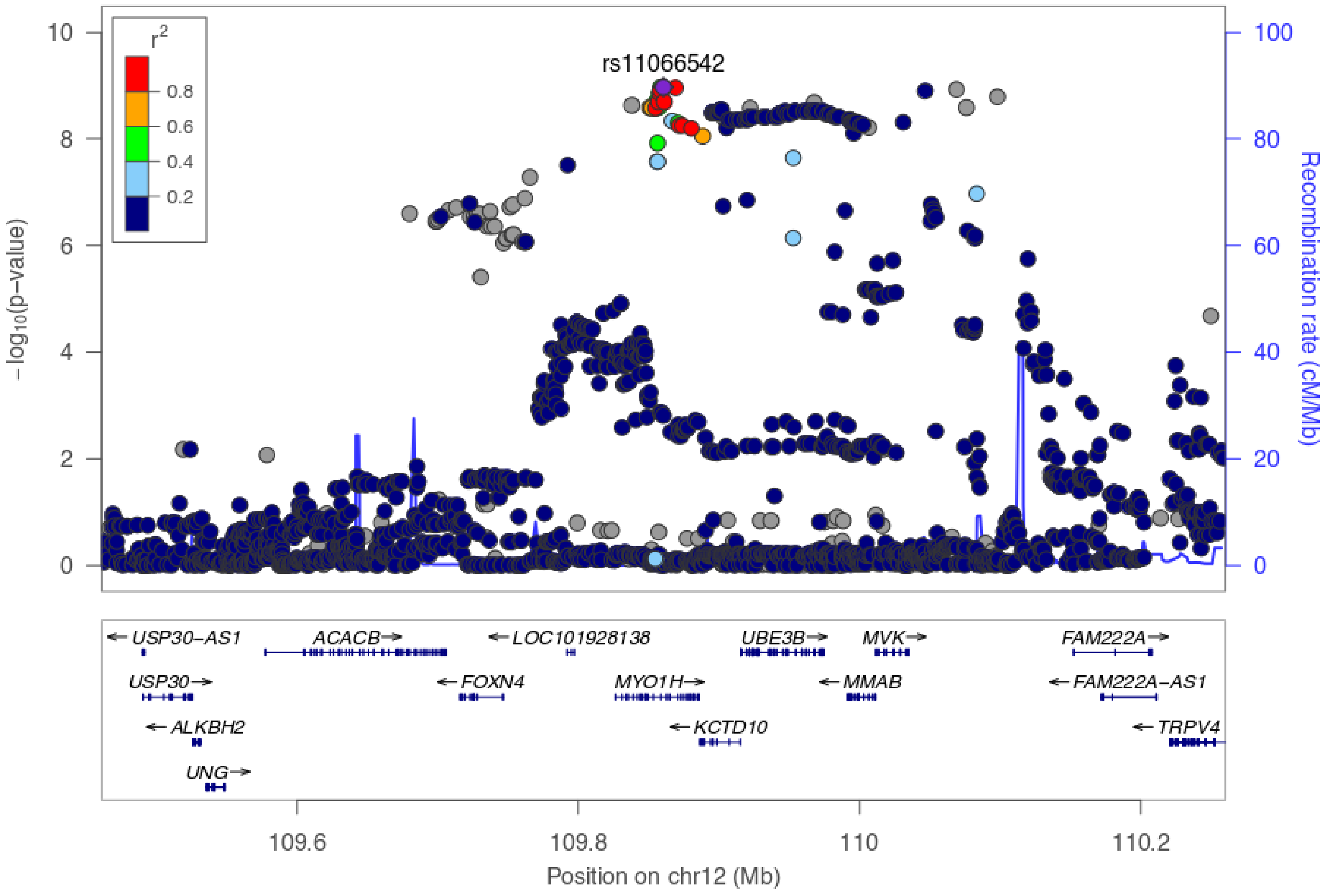

r2
0.8
0.6
0.4
0.2
rs11066542
−log10(p−value)
Recombination rate (cM/Mb)
10
8
6
4
2
0
100
80
60
40
20
0
USP30−AS1
ACACB
LOC101928138
UBE3B
MVK
FAM222A
USP30
FOXN4
MYO1H
MMAB
FAM222A−AS1
ALKBH2
KCTD10
TRPV4
UNG
109.6
109.8
110
110.2
Position on chr12 (Mb)

E

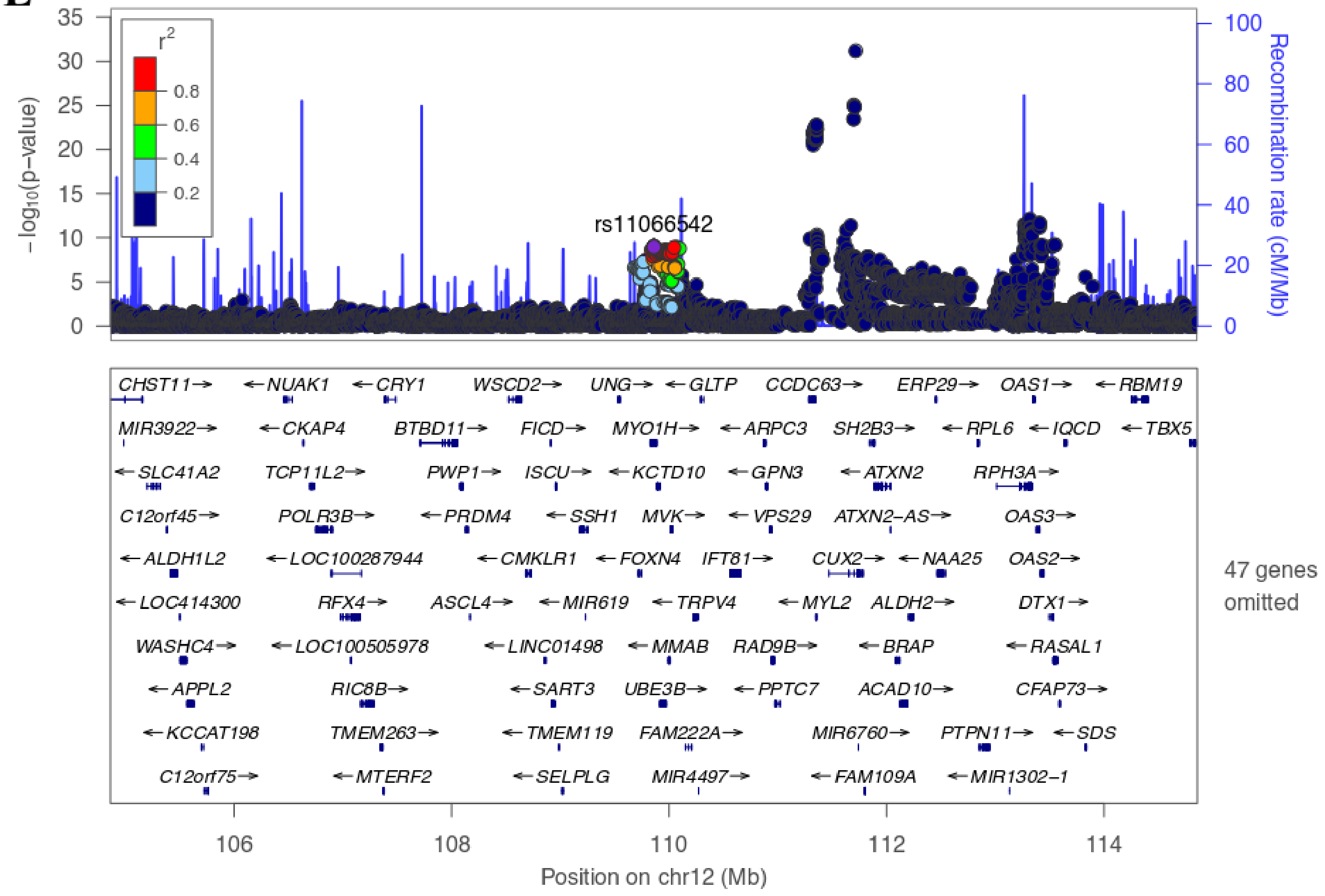

35
30
25
20
15
10
5
0
r2
0.8
0.6
0.4
0.2
−log10(p−value)
rs11066542
100
80
60
40
20
0
Recombination rate (cM/Mb)
CHST11→
←NUAK1
←CRY1
WSCD2→
UNG→
←GLTP
CCDC63→
ERP29→
OAS1→
←RBM19
MIR3922→
←CKAP4
BTBD11→
FICD→
MYO1H→
←ARPC3
SH2B3→
←RPL6
←IQCD
←TBX5
←SLC41A2
TCP11L2→
PWP1→
ISCU→
←KCTD10
←GPN3
←ATXN2
RPH3A→
C12orf45→
POLR3B→
←PRDM4
←SSH1
MVK→
←VPS29
ATXN2−AS→
OAS3→
←ALDH1L2
←LOC100287944
←CMKLR1
←FOXN4
IFT81→
CUX2→
←NAA25
OAS2→
←LOC414300
RFX4→
ASCL4→
←MIR619
←TRPV4
←MYL2
ALDH2→
DTX1→
WASHC4→
←LOC100505978
←LINC01498
←MMAB
RAD9B→
←BRAP
←RASAL1
←APPL2
RIC8B→
←SART3
UBE3B→
←PPTC7
ACAD10→
CFAP73→
←KCCAT198
TMEM263→
←TMEM119
FAM222A→
MIR6760→
PTPN11→
←SDS
C12orf75→
←MTERF2
←SELPLG
MIR4497→
←FAM109A
←MIR1302−1
47 genes omitted
106
108
110
112
114
Position on chr12 (Mb)

F

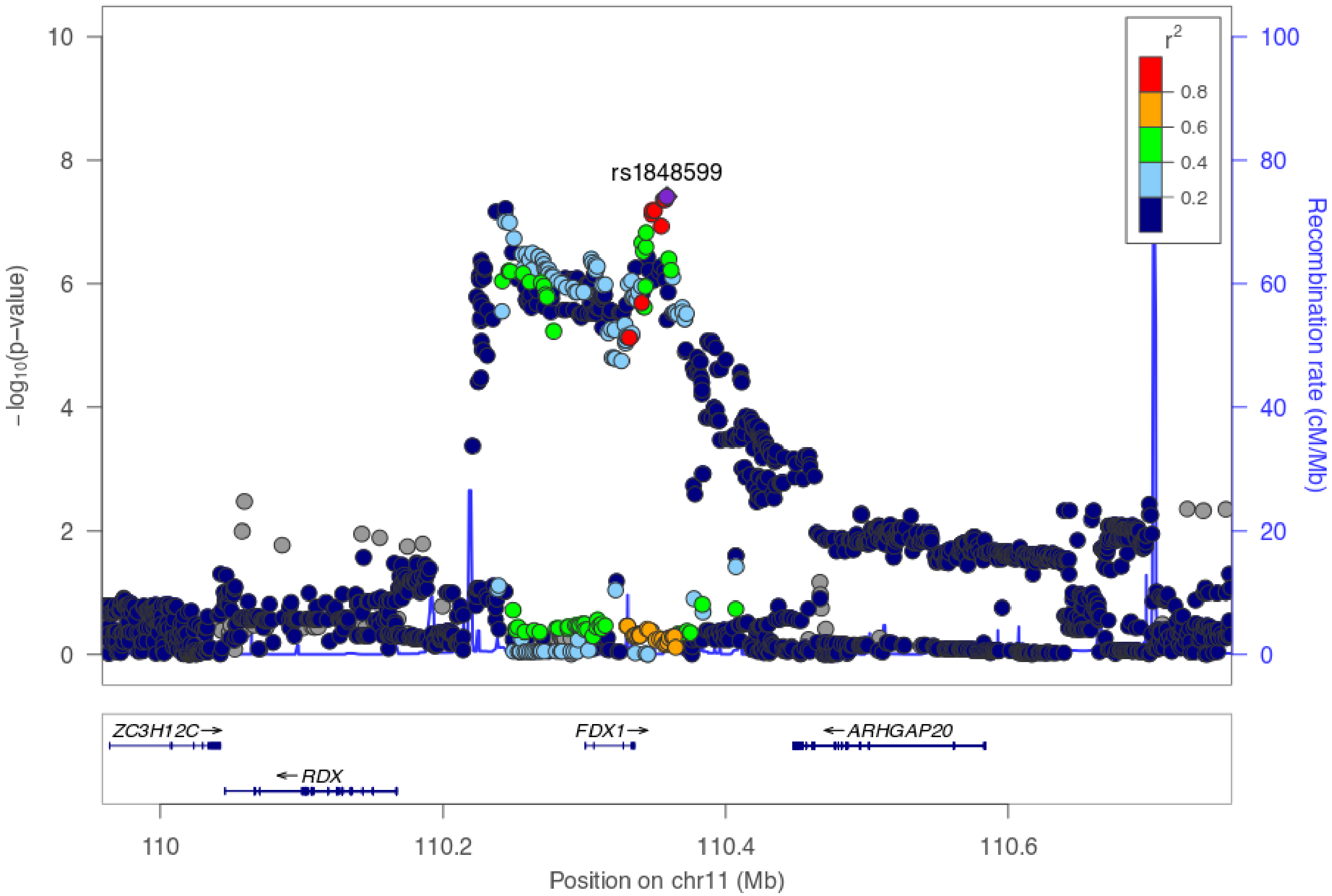
rs1848599
r2
0.8
0.6
0.4
0.2
10
8
6
4
2
0
−log10(p−value)
100
80
60
40
20
0
Recombination rate (cM/Mb)
ZC3H12C→
FDX1→
←ARHGAP20
←RDX
110
110.2
110.4
110.6
Position on chr11 (Mb)

G

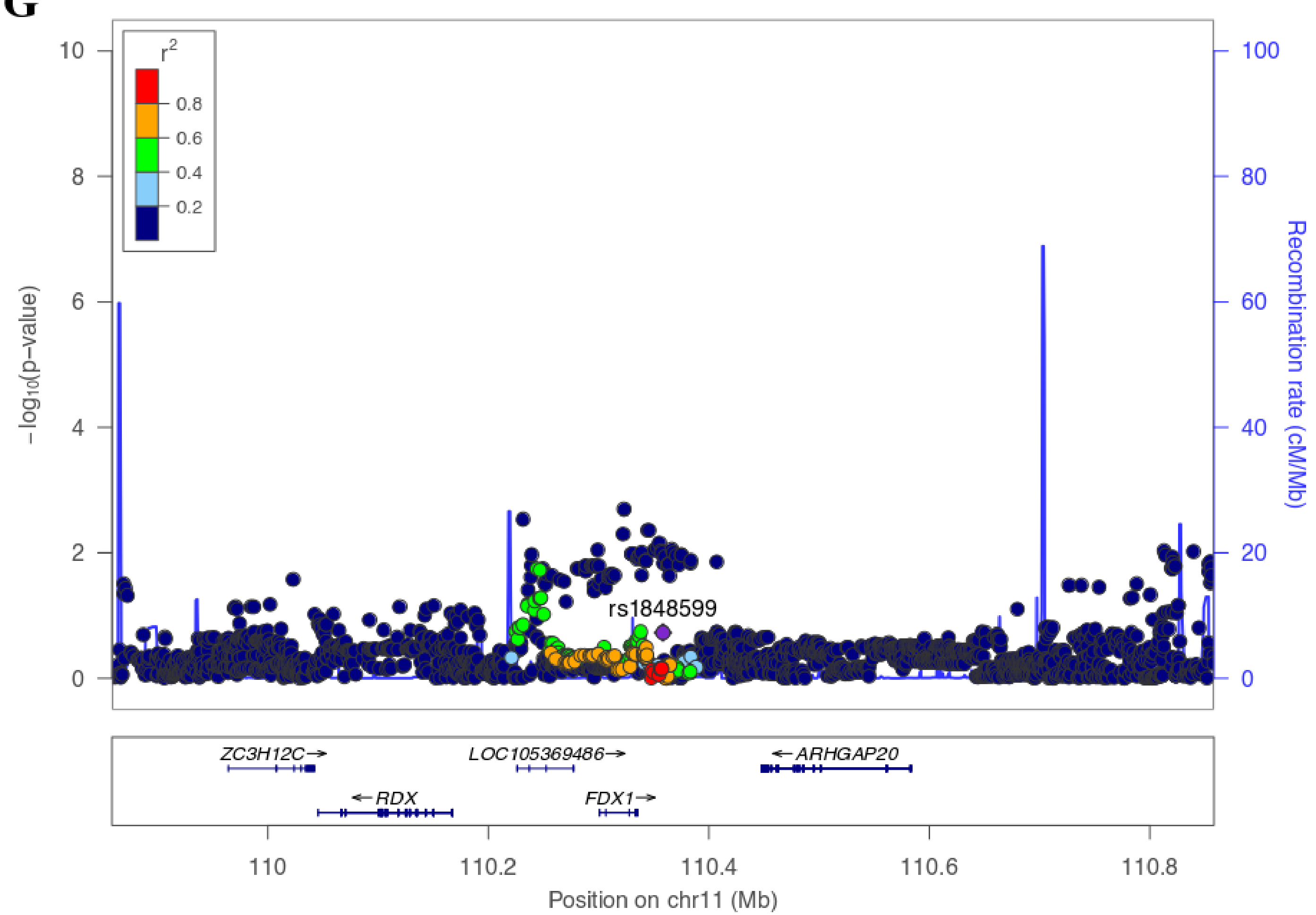


**Supplemental Figure 4. Regional plots for novel loci**

The *x* axis denotes chromosomal location and the *y* axis denotes -$\log_{10}$ *P* value for each SNP. Lead variant is shown in purple. Variants in linkage disequilibrium with the lead variant are shown in red ($r^2 > 0.8$), orange ($r^2 > 0.6$), green ($r^2 > 0.4$), and light blue ($r^2 > 0.2$). **A**, 1q21 (rs6587520). **B**, 10q26 (rs2257129). **C**, 11q22 (rs10488763). **D**, 12q24 (rs11066542). **E**, 12q24 (rs11066542) conditioning on rs79105258 (a proxy for *BRAP-ALDH2*). **F**,

11q22 (rs1848599). **G**, 11q22 (rs1848599) conditioning on rs10488763. Linkage disequilibrium was calculated based on 1000 Genomes Project phase 1 European samples (A, B, and C) and East Asian samples (D, E, F and G).

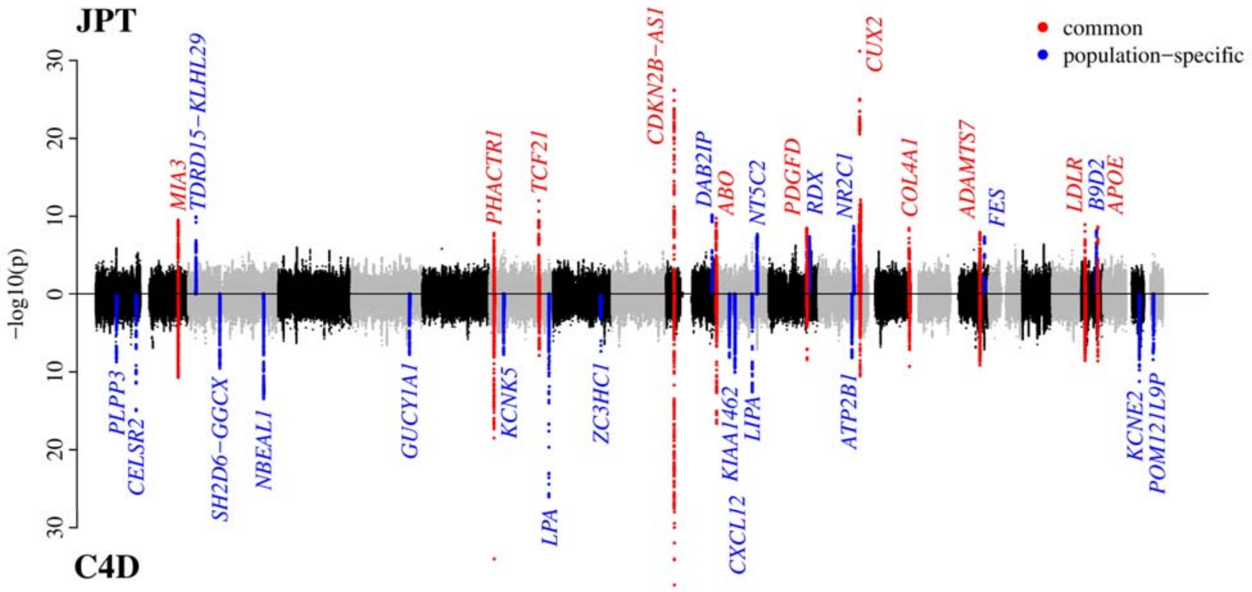


**Supplementary Figure 5. Miami plot of myocardial infarction in the Japanese meta-analysis and the CARDIoGRAMplusC4D 1000 Genomes-based GWAS**

The upper panel shows the result of the Japanese meta-analysis, while the bottom panel presents the myocardial infarction subphenotype meta-analysis result in the CARDIoGRAMplusC4D 1000 Genomes-based GWAS. The *x* axis denotes chromosomal location and the *y* axis denotes $-\log_{10}$ *P* value for each SNP. Red and blue dots show common and population-specific genome-wide significant loci, respectively.

JPT, Japanese; C4D, CARDIoGRAMplusC4D.
